# Mechanisms of spontaneous Ca^2+^ release-mediated arrhythmia in a novel 3D human atrial myocyte model: II. Ca^2+^-handling protein variation

**DOI:** 10.1101/2022.07.15.500278

**Authors:** X. Zhang, C.E.R. Smith, S. Morotti, A.G. Edwards, D. Sato, W.E. Louch, H. Ni, E. Grandi

**Affiliations:** Department of Pharmacology, University of California, Davis, CA 95616, USA; University of Oslo, Norway; Simula Research Laboratory, Oslo, Norway

**Keywords:** Atria, Calcium, Arrhythmia, Transverse-axial tubule, Calcium-handling proteins

## Abstract

Disruption of the transverse-axial tubule system (TATS) in diseases such as heart failure and atrial fibrillation occurs in combination with changes in the expression and distribution of key Ca^2+^- handling proteins. Together this ultrastructural and ionic remodeling is associated with aberrant Ca^2+^ cycling and electrophysiological instabilities that underly arrhythmic activity. However, due to the concurrent changes in TATs and Ca^2+^-handling protein expression and localization that occur in disease it is difficult to distinguish their individual contributions to the arrhythmogenic state. To investigate this, we applied our novel 3D human atrial myocyte model with spatially detailed Ca^2+^ diffusion and TATS to investigate the isolated and interactive effects of changes in expression and localization of key Ca^2+^-handling proteins and variable TATS density on Ca^2+^- handling abnormality driven membrane instabilities. We show that modulating the expression and distribution of the sodium-calcium exchanger, ryanodine receptors, and the sarcoplasmic reticulum (SR) Ca^2+^ buffer calsequestrin have varying pro and anti-arrhythmic effects depending on the balance of opposing influences on SR Ca^2+^ leak-load and Ca^2+^-voltage relationships. Interestingly, the impact of protein remodeling on Ca^2+^-driven proarrhythmic behavior varied dramatically depending on TATS density, with intermediately tubulated cells being more severely affected compared to detubulated and densely tubulated myocytes. This work provides novel mechanistic insight into the distinct and interactive consequences of TATS and Ca^2+^-handling protein remodeling that underlies dysfunctional Ca^2+^ cycling and electrophysiological instability in disease.

**Key Points:**

- In our companion paper we developed a 3D human atrial myocyte model, coupling electrophysiology and Ca^2+^ handling with subcellular spatial details governed by the transverse-axial tubule system (TATS).
- Here we utilize this model to mechanistically examine the impact of TATS loss and changes in the expression and distribution of key Ca^2+^-handling proteins known to be remodeled in disease on Ca^2+^ homeostasis and electrophysiological stability.
- We demonstrate that varying the expression and localization of these proteins has variable pro- and anti-arrhythmic effects with outcomes displaying dependence on TATS density.
- Whereas detubulated myocytes typically appear unaffected and densely tubulated cells seem protected, the arrhythmogenic effects of Ca^2+^ handling protein remodeling are profound in intermediately tubulated cells.
- Our work shows the interaction between TATS and Ca^2+^-handling protein remodeling that underlies the Ca^2+^-driven proarrhythmic behavior observed in AF and may help to predict the effects of antiarrhythmic strategies at varying stages of ultrastructural remodeling.

## Introduction

It is widely accepted that Ca^2+^ abnormalities can precipitate atrial fibrillation (AF), the most common cardiac arrhythmia. Both ionic remodeling and ultrastructural changes have been associated with aberrant Ca^2+^ signaling, excitation-contraction coupling, and AF (Nattel & Dobrev, 2012; Nattel & Harada, 2014; Dobrev & Wehrens, 2017; Denham *et al*., 2018). Disruption of the transverse-axial tubular system (TATS) is one hallmark of AF structural remodeling that likely occurs early in disease progression and contributes to the dysfunction observed. In a companion paper, we developed a three-dimensional model of the human atrial myocyte coupling electrophysiology, whole-cell and local Ca^2+^ handling, and subcellular ultrastructural details, to interrogate the mechanisms by which TATS variability and loss affect human atrial physiology (Zhang et al.). Our simulation predicted that TATS loss *per se* enhances vulnerability to proarrhythmic behaviors (i.e., spontaneous calcium releases, SCRs, delayed afterdepolarizations, DADs) by altering subcellular Ca^2+^ signaling. Nevertheless, the reduced density and regularity of the TATS in disease is accompanied by altered channel and transporter expression and function, as well as subcellular redistribution of ion channels, transporters, and Ca^2+^ handling proteins.

Well established hallmarks of ionic remodeling in AF include reduced L-type Ca^2+^ channel (LCC) current (Lenaerts *et al*., 2009), increased inward rectifier K^+^ current (I_K1_) and constitutively activated acetylcholine-activated K^+^ current (I_K,ACh_) (Bosch *et al*., 1999), enhanced Na^+^/Ca^2+^ exchanger (NCX) activity (Lenaerts *et al*., 2009; Voigt *et al*., 2012), and hyperphosphorylated ryanodine receptors (RyRs) (Vest *et al*., 2005; Neef *et al*., 2010; Voigt *et al*., 2012). The basal subcellular localization of these proteins in the atria, and whether they are altered in disease, in not well understood. LCCs distribute equally in tubules and crest areas of the sarcolemma in control human atrial myocytes (Glukhov *et al*., 2015). Given atrial LCC current amplitude likely depends on TATS density as shown in rat ventricular myocyte experiments (Frisk *et al*., 2014), TATS loss in chronic AF might contribute to the observed reduction in current density. Recently, in human atrial myocytes from both normal sinus rhythm and chronic AF patients, RyR and the sarcoplasmic reticulum (SR) Ca^2+^ buffer calsequestrin (CSQ) were found more densely expressed near the cell periphery vs. the cell interior (Herraiz-Martínez *et al*., 2022). This is similar to what has previously been shown in rat atrial myocytes, where RyR-CSQ colocalization was reduced in the cell interior vs. periphery (Schulson *et al*., 2011). To our knowledge, NCX localization in human atrial myocytes has not yet been investigated, and while NCX is expressed at both the cell surface and in the TATS membrane, the precise quantitative distribution may vary in different species and cardiac regions (Melnyk *et al*., 2005; Scriven *et al*., 2010; Schulson *et al*., 2011).

The available data on ion channels and Ca^2+^ handling protein expression, localization, and function in human atrial myocytes is limited and may not be representative of the heterogeneous human atrial myocyte population in tissue. There are known regional differences across the atria, e.g., in right vs. left atrium (Arora *et al*., 2017), pulmonary vein vs. free atrial wall (Melnyk *et al*., 2005), and insight from isolated cells is also complicated by TATS damage from the enzymatic digestion (Chen *et al*., 2015). Furthermore, it remains unclear whether ionic and ultrastructural remodeling interacts to affect arrhythmogenic propensity. Indeed, because Ca^2+^ handling protein expression, localization, and regulatory states change simultaneously with TATs in disease (Brandenburg *et al*., 2016; Yue *et al*., 2017), their effects cannot easily be separated in experiments. To address this, we utilized our new 3D model of the human atrial myocyte with spatially detailed Ca^2+^ diffusion and TATS (Zhang et al.) to investigate the isolated and interactive effects of altered TATS and changes in expression and localization of key Ca^2+^-handling proteins (i.e., NCX, RyR, and CSQ) on Ca^2+^-driven membrane instabilities. We focused on these proteins because both 1) NCX and RyR are altered in AF and emerged as key mediators of TATS loss-induced diastolic instabilities in Ca^2+^ and membrane voltage (V_m_), and 2) RyR and its regulator CSQ have been found to distribute non-uniformly between the cell periphery and interior. In this study, we found that TATS loss and Ca^2+^ handling protein remodeling collectively promote arrhythmia. Cells with intermediate TATS density were most sensitive to changes in protein expression and localization, whereby ultrastructural and Ca^2+^ handling remodeling synergistically contributed to the proarrhythmic outcomes. Conversely, when the TATS is depleted, changes in protein expression and distribution have little effect. Our study demonstrates the interactive contributions of TATS and Ca^2+^-handling protein expression and distribution on maintaining Ca^2+^ and membrane potential stability in human atrial myocytes and provides novel model-based mechanistic insight that may guide future therapeutic anti-AF strategies.

## Methods

We simulated human atrial myocyte electrophysiology and Ca^2+^ handling using our recently developed model, integrating transmembrane voltage dynamics, spatially-detailed Ca^2+^ diffusion, and varying TATS structure (Zhang et al.). We modified the expression and subcellular distribution of various Ca^2+^ handling proteins (i.e., namely NCX, RyR, and CSQ), as detailed below. To investigate the interactive effects of altering protein expression and localization and varying tubular structures, we conducted simulations in cells with dense, median, and sparse TATS taken from the population of TATS structures described in our companion paper (Zhang et al.).

### Varying subcellular distribution of NCX, RyR, and CSQ

To investigate how the heterogeneous subcellular localization of NCX, RyR, and CSQ impact electrophysiology and Ca^2+^ handling, we systematically varied the relative density of NCX, RyR or CSQ localized to surface Ca^2+^ release units (CRUs, i.e., the 2 outer CRU layers near cell surface) vs. those localized to inner (i.e., non-surface) CRUs while keeping the whole-cell total expression unchanged (**Fig. 1**). Specifically, we varied the ratio of surface/inner CRU protein density between 0.5 and 2 to reflect the surface-to-central gradient in the subcellular distribution of RyR and CSQ observed in experiments (Herraiz-Martínez *et al*., 2022).

**Figure 1.**
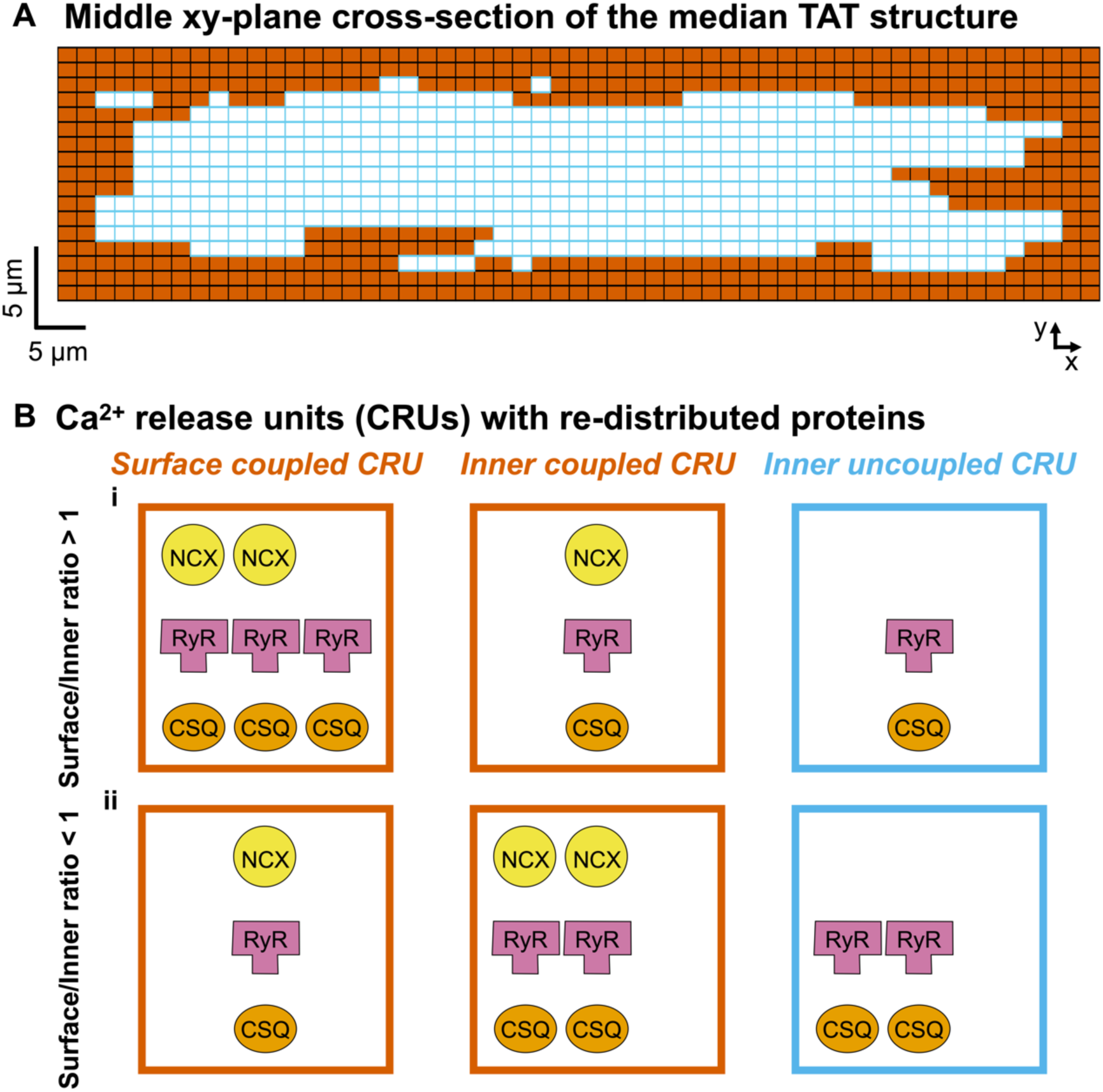
Varying Ca^2+^ handling proteins (Na^+^-Ca^2+^ exchanger, NCX, Ryanodine receptor, RyR, and calsequestrin, CSQ) distribution. **A)** Two-dimensional section of a middle XY plane in the intermediate TAT structure shown in the companion paper (Zhang et al.). Surface coupled CRUs and inner coupled CRUs are in orange, and uncoupled CRUs are in light blue. **B)** Schematic of surface coupled (left), central coupled (middle), and uncoupled (right) CRUs when surface/inner CRU protein expression ratio > 1 (**i**) or < 1 (**ii**). When varying the NCX (yellow) distribution, we scale the NCX current density in inner coupled and surface coupled CRUs accordingly. When varying the RyR distribution, we scale the relative expression of RyRs (pink) in the inner (coupled and uncoupled) CRUs vs. surface coupled CRUs. When varying the CSQ (orange) distribution, we scale the relative CSQ expression in the inner (coupled and uncoupled) CRUs vs. surface coupled CRUs.

### Varying the whole-cell expression of NCX, RyR, and CSQ

The whole-cell expression (i.e., NCX maximal transport rate, number of RyRs per CRU, and CSQ maximum buffering capacity, B_CSQ_) was scaled to explore the effects of experimentally-determined variation ranges of NCX (Schotten *et al*., 2002; El-Armouche *et al*., 2006; Lenaerts *et al*., 2009; Voigt *et al*., 2012; Yue *et al*., 2017), RyR (Ohkusa *et al*., 1999; Neef *et al*., 2010; Voigt *et al*., 2014; Greiser *et al*., 2014), and CSQ (Lüss *et al*., 1999; Faggioni *et al*., 2014). Specifically, NCX was varied between 10% and 200%, RyR was varied between 10% and 140%, and CSQ was varied between 50% and 140% of the baseline expression outlined in Zhang et al.

Since CSQ is not only a luminal SR Ca^2+^ buffer but also a regulator of RyR function (Terentyev *et al*., 2003; Györke *et al*., 2004; Knollmann *et al*., 2006; Restrepo *et al*., 2008), to dissect the relative and combined contributions of these processes we investigated three scenarios when performing simulations to study the impacts of varying CSQ: 1) only luminal SR Ca^2+^ buffering is affected; 2) only RyR regulation is affected; or 3) both SR buffering and RyR regulation are affected. To assess this, changes in Ca^2+^ buffering CSQ were implemented by modifying the B_CSQ_ in the equations of junctional SR (JSR) Ca^2+^ rapid equilibrium approximation, i.e., β([Ca^2+^]_JSR_) in the companion paper by Zhang et al; whereas changes in RyR gating were implemented by modifying the B_CSQ_ in RyR luminal-Ca^2+^-dependent transient rate equations, i.e., k_14_ and k_23_ in Table 5 of the companion paper by Zhang et al.

### Numerical methods and simulation protocols

Simulation details are as in the companion paper (Zhang et al.). Briefly, our human atrial myocyte model was implemented in C++ and parallelized using OpenMP 5.1 (Dagum & Menon, 1998). The ordinary differential equations (ODEs) and subcellular Ca^2+^ diffusion were solved using an explicit Forward Euler method except that LCC current (I_Ca_) and RyR gating behaviors were described stochastically as in (Restrepo *et al*., 2008; Sato & Bers, 2011), and that ODEs of fast Na^+^ current (I_Na_) were solved using the Rush-Larsen scheme (Rush & Larsen, 1978). The time step was 0.01 ms.

### Measurement of Ca^2+^-handling abnormalities, V_m_ instabilities and differences in Ca^2+^ release between CRUs

To compare Ca^2+^ signaling in surface CRUs, inner coupled CRUs (those CRUs located away from the cell surface but coupled to tubules), and inner uncoupled CRUs, the [Ca^2+^]_Cyto_ within each group of CRUs throughout the cell was averaged. Ca^2+^-handling abnormalities and V_m_ instabilities associated with changes in protein expression and distribution were examined for SCRs, DADs, and spontaneous action potentials (SAPs). Detailed descriptions of SCR and DAD measurements are provided in our companion paper (Zhang et al.). Briefly, cells were paced to steady state with the stimulation then stopped and the subsequent diastolic period used for analysis in MATLAB R2021. SCRs were defined as Ca^2+^-release events with an amplitude > 0.3 µM, with DADs defined as membrane depolarizations > 10 mV. SAPs were deemed present when the amplitude of a DAD exceeded 70 mV.

### Measurements of alternans

To understand the effects of varying expression and distribution of Ca^2+^ handling proteins on the regularity of Ca^2+^ release, the occurrence of pacing-induced alternans in [Ca^2+^]_Cyto_ and V_m_ traces were analyzed as done in experiments (Myles *et al*., 2011; Hammer *et al*., 2015). Specifically, Ca^2+^ alternans were distinguished when the calculated index (r) that compares the alternating averaged smaller (S) and larger (L) Ca^2+^ transient amplitudes (i.e., r = 1 – S/L) was over 0.08. APD alternans were identified when the difference between the averaged longer APD_90_ and the averaged shorter APD_90_ (ΔAPD_90_) was over 10 ms.

### Source code

The source code of our new 3D human atrial cell model can be accessed from http://elegrandi.wixsite.com/grandilab/downloads and https://github.com/drgrandilab.

## Results

### Lowering NCX promotes SCRs, but has biphasic effects on DADs and SAPs, depending on the balance between increased SCRs and reduced ΔV_m_/ΔCa^2+^ gain

To reveal how NCX expression and TATS changes affect arrhythmic biomarkers (i.e., SCRs, DADs, and SAPs), we varied NCX expression in the human atrial myocyte models with sparse, intermediate, and dense TATS. We found that the SCR rate threshold monotonically decreases with reduced NCX expression and increased NCX associated with fewer SCRs, especially in cells with sparse TATS (**Fig. 2Ai**). While low NCX expression is associated with decreased SCR rate threshold, it conversely increases the rate thresholds of both DADs and SAPs when below 20%-40% of its baseline value. Interestingly, at low expression levels above 40% of baseline, DAD and SAP rate threshold is decreased matching SCR-NCX expression dependence (**Fig. 2Aii-iii**). This is demonstrated in the representative voltage (**i**) and [Ca^2+^]_Cyto_ (**ii**) traces in (**Fig. 2B**) where the biphasic effects of NCX reduction on DADs and SAPs (but not SCRs) are highlighted by the occurrence of SCRs at 0.2 and 0.5 NCX expression but DADs and SAPs solely occurring at 0.5. This result suggests that [Ca^2+^]_Cyto_-V_m_ coupling changes with low vs. high NCX expression. Interestingly, while the SCR pacing threshold is lower in cells with sparser TATS, SAPs occur at lower pacing rates in cells with intermediate and dense TATS, suggesting that NCX expression changes have the greatest impact on [Ca^2+^]_Cyto_-V_m_ coupling in cells with relatively intact TATS. We further analyzed the model to understand the mechanism by which reducing NCX promotes SCRs but has biphasic effects on DADs and SAPs. We found that lowering NCX expression reduced NCX Ca^2+^ extrusion (**Fig. 2Ci**), leading to elevated diastolic cleft Ca^2+^ concentration, augmented diastolic RyR open probability (P_O_) and leak, and increased SCRs and DADs. This mechanism resembles that suggested to be responsible for elevated SCR and DAD propensity in cells with sparse TATS in our companion paper (Zhang et al.). Conversely, less NCX reduced SCR-induced ΔV_m_, especially in cells with intermediate and dense TATS (**Fig. 2Cii**), leading to decoupling between SCR and DAD/SAPs and subsequent biphasic changes of DADs and SAPs.

**Figure 2.**
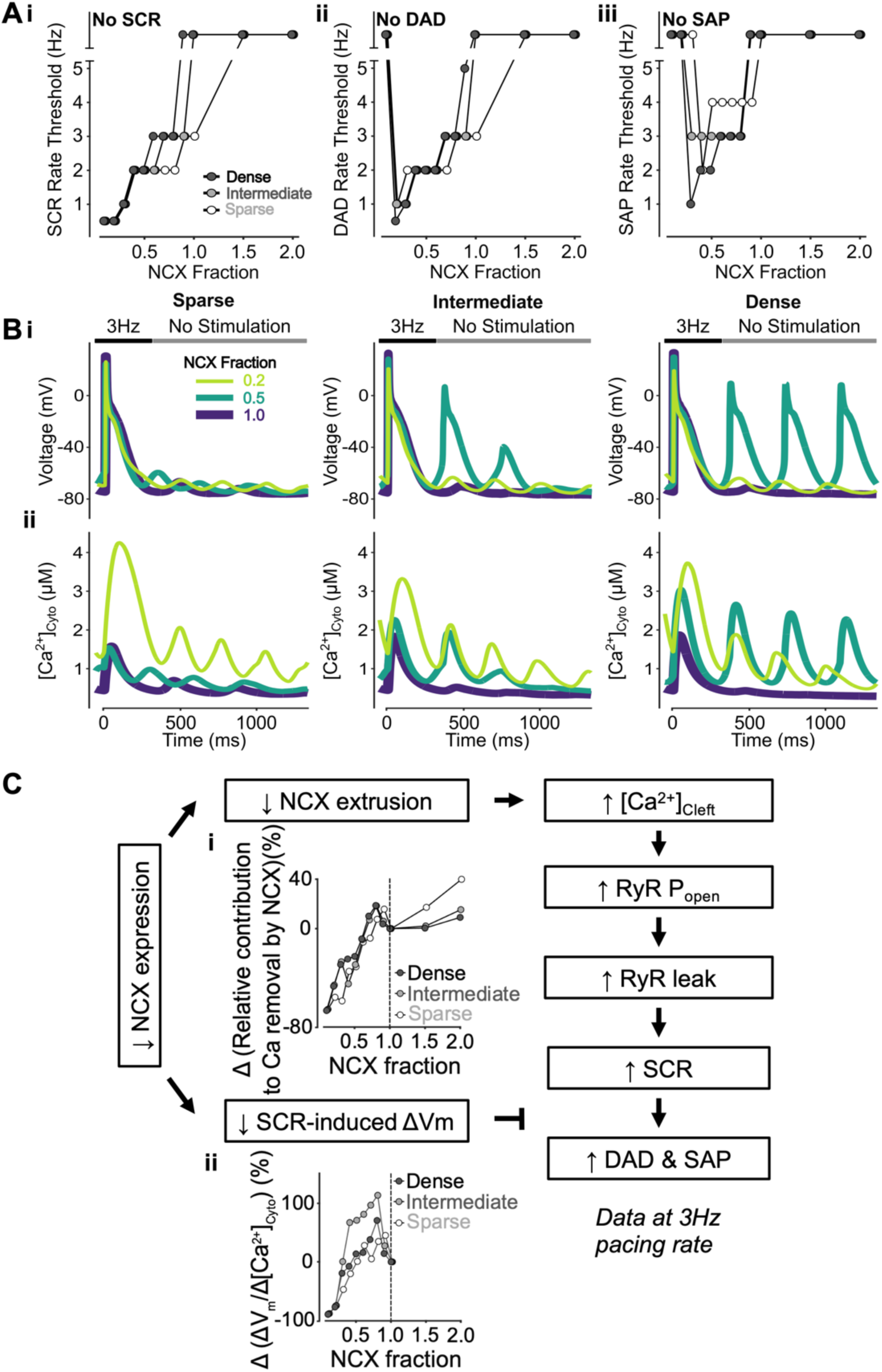
Inhibition of NCX promotes SCRs but has biphasic effects on DADs and spontaneous APs (SAP). **A)** Reducing NCX monotonically decreases the rate threshold of SCRs (i), whereas the rate thresholds of DADs (ii) and SAP, defined as DADs with amplitudes over 70 mV, (iii) display biphasic dependence. **B)** Effect of altered NCX fraction on voltage (i) and global cytosolic Ca^2+^ concentration (ii) in cells with sparse (left), intermediate (middle), and dense (right) tubules following pacing at 3 Hz to examine the occurrence of DADs, SAPs and SCRs. **C)** Mechanism underlying NCX inhibition promoting SCRs but having biphasic effects on DADs and SAP. Biomarkers were determined from the first 100 ms of no-stimulation period and normalized to those of cells with a retained NCX fraction of 1.0. Lower NCX expression is associated with reduced NCX contribution to Ca^2+^ extrusion (i) and smaller SCR-induced voltage changes/SCR amplitude ratio (ii). Less NCX extrusion results in elevated cleft Ca^2+^ concentration, augmented RyR P_o_ and leak leading to increased SCRs. While SCRs lead to DADs and SAP this transition is abated by the reduced SCR-induced changes in V_m_, thus explaining the biphasic effects of reduced NCX.

We further sought to determine whether regional variations exist in the effects of varying NCX expression and TATS density on the pacing threshold for SCR. To do so, we analyzed the averaged Ca^2+^ transient of surface CRUs, inner CRUs coupled to TATS (inner coupled), and inner CRUs that are not coupled to TATS (inner uncoupled) in each cell in the population with varying NCX expression and TATS density and quantified the biomarkers of local Ca^2+^ transient. As expected, reduced NCX lowered the rate threshold for SCRs in all CRUs (**Fig. 3Ai**). The representative traces of average cytosolic Ca^2+^ concentration following pacing at 3 Hz in cells with dense (**Fig. 3Bi**), intermediate (**Fig. 3Bii**), and sparse TATS (**Fig. 3Biii**) also show larger SCRs with lower NCX expression. Interestingly, when NCX was severely reduced (scaling factor of 0.2 of baseline expression), Ca^2+^ alternans were observed in all CRUs from cells with intermediate and dense TATS (**Fig. 3Bi-ii**) indicating enhanced arrhythmic effect of lower NCX expression in tubulated cells. Overall, our simulations suggest that NCX inhibition shifts the balance between increased SCRs and reduced ΔV_m_/ΔCa^2+^ gain. While cells with sparse TATS consistently display a lower SCR pacing threshold, when SCRs occur in more densely tubulated cells they are greater in magnitude and the SAP pacing threshold appears more sensitive to variations in NCX levels compared to sparsely tubulated cells.

**Figure 3.**
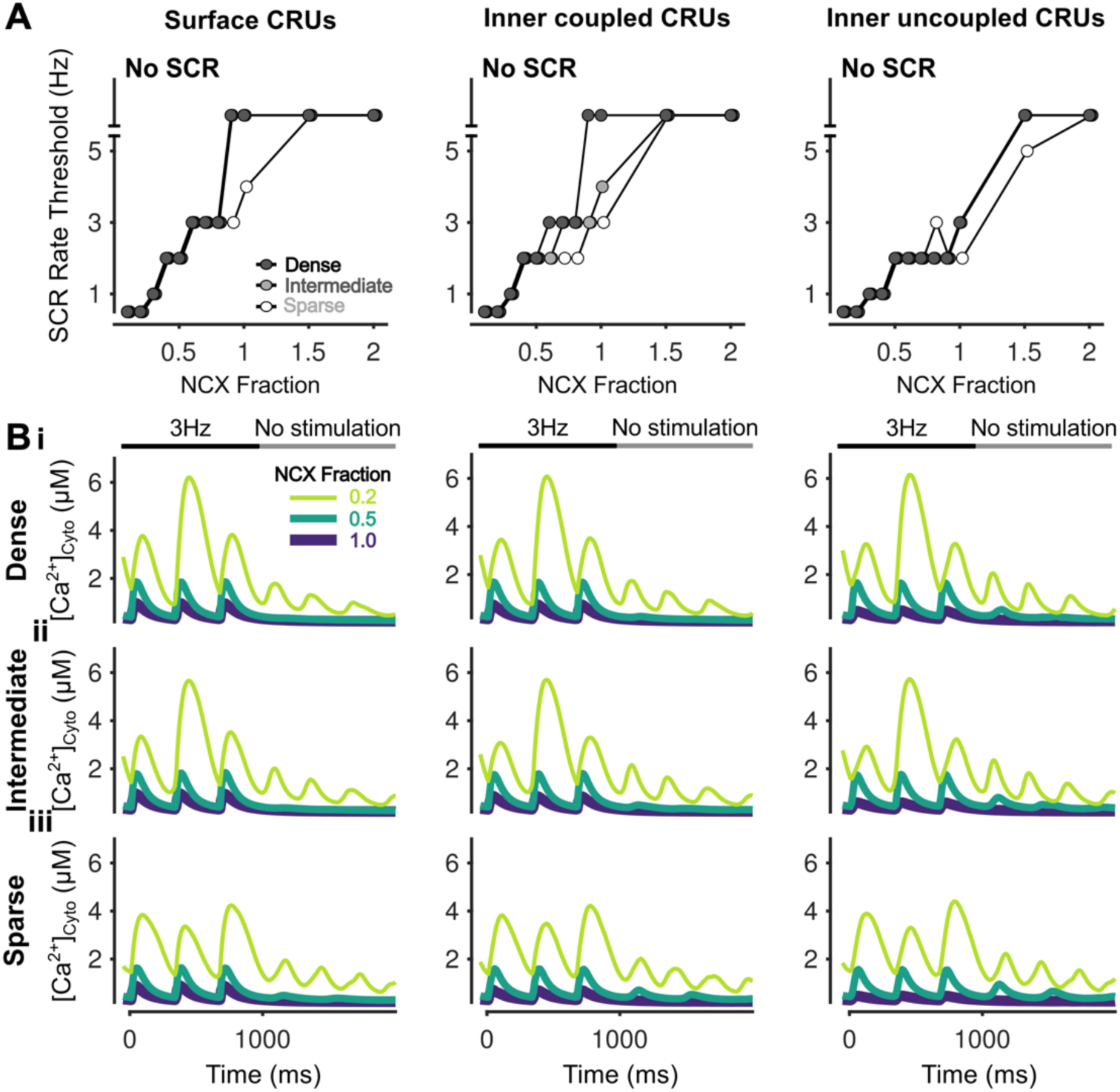
Inhibition of NCX promotes simultaneous SCRs in all CRUs. **A)** Rate threshold for local SCRs of surface (left), inner coupled (middle) and inner uncoupled (right) CRUs in dense, intermediate and sparsely tubulated cells. **B)** Cytosolic Ca^2+^ concentration of surface (left), inner coupled (middle) and inner uncoupled (right) CRUs from cells with dense (i), intermediate (ii), and sparse (iii) tubules following pacing at 3 Hz showing SCRs and alternans for all CRUs at NCX Fraction of 0.2.

### Increasing surface/inner CRU NCX expression ratio promotes SCRs and DADs by elevating inner [Ca^2+^]_Cleft_ and RyR leak

After characterizing the impact of changes to global NCX expression on SCRs, DAD, and SAPs, we sought to investigate the effects of varying the spatial distribution of NCX and TATS density. To do this we altered the ratio of NCX in surface/inner CRUs in the human atrial myocyte models with sparse, intermediate, and dense TATS. Our simulations indicate that increasing the ratio of surface to inner CRU NCX expression decreases the rate threshold of SCRs (**Fig. 4Ai**) and DADs (**Fig. 4Aii**), without triggering SAPs (**Fig. 4Aiii**). This effect is more pronounced in cells with dense and intermediate TATS (**Fig. 4Ai-ii**), as also shown in the representative traces of V_m_ (**Fig. 4Bi**) and [Ca^2+^]_Cyto_ (**Fig. 2Bii**). We found that increasing the surface-to-inner CRU NCX expression ratio decreased inner NCX Ca^2+^ extrusion but enhanced NCX Ca^2+^ extrusion at the surface. While [Ca^2+^]_Cleft_ in surface CRUs remained unaffected (**Fig. 4Ci**), lower NCX expression in the inner CRUs lead to increased [Ca^2+^]_Cleft_ in both inner coupled (**Fig. 4Cii**) and uncoupled (**Fig. 4Ciii**) CRUs, thus causing elevated RyR P_O_ and leak, which promotes SCRs and DADs. The consequences of heterogeneous NCX distribution were especially pronounced in inner coupled CRUs in cells with dense and intermediate TATS (**Fig. 5A-B**). Overall, we found that increasing the surface/inner CRU NCX expression ratio promotes SCRs, especially in inner coupled CRUs, and DADs, but does not affect SAPs. These effects are marked in cells with intermediate and dense TATS, suggesting that loss of TATS and reduced NCX expression in inner CRUs favors SCRs and DADs via a similar mechanism, i.e., the elevation of inner [Ca^2+^]_Cleft_ and RyR leak.

**Figure 4.**
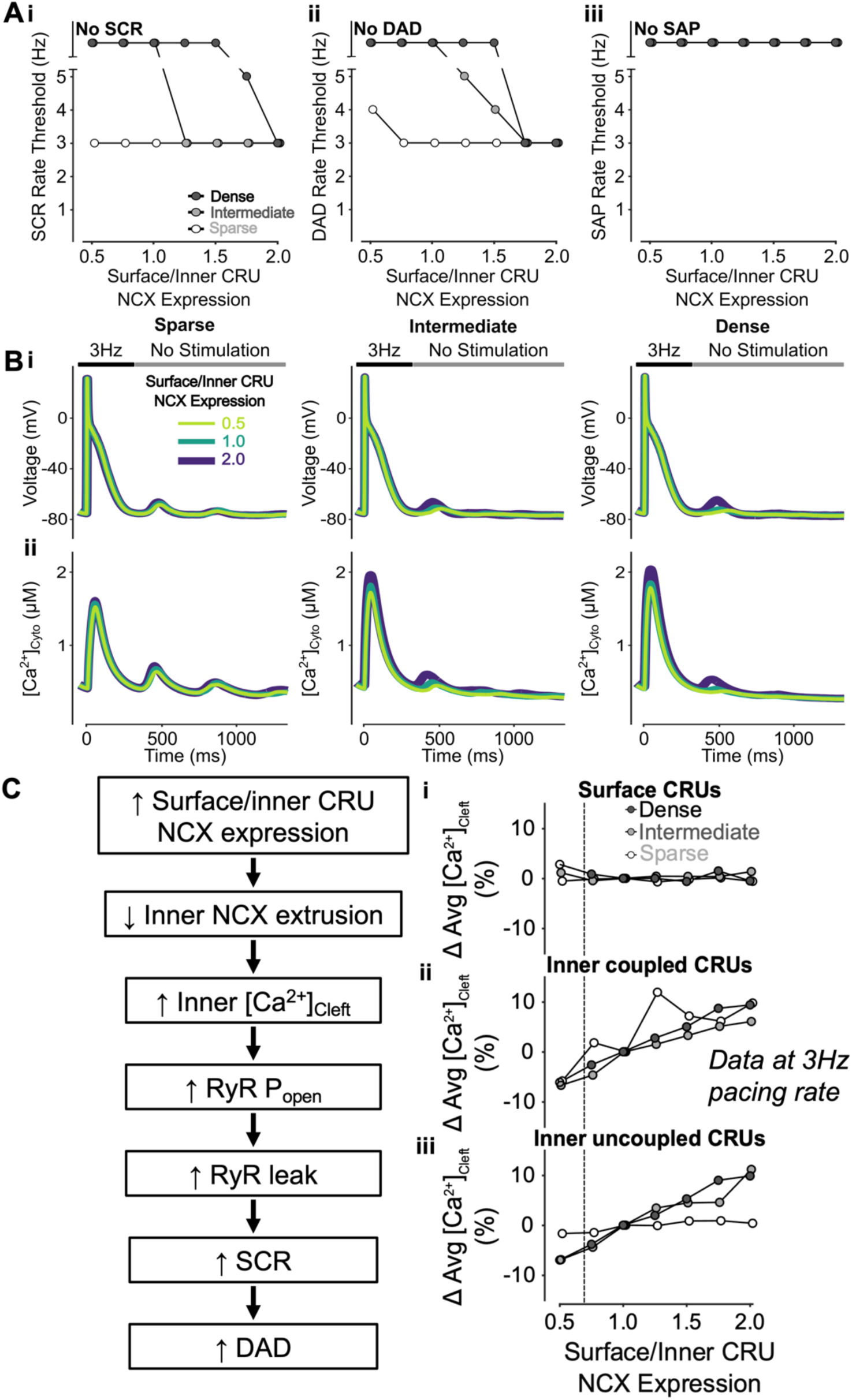
Increasing surface/inner CRU NCX expression ratio inhibits SCRs and DADs but does not affect SAPs. **A)** Increasing surface/inner CRU NCX expression ratio (i.e., increasing NCX density in surface CRUs without changing whole-cell NCX expression) monotonically decreases the rate threshold of SCRs (i) and DADs (ii), whereas SAP remains absent in all conditions. This effect is greater in cells with dense and intermediate tubular structures vs cells with sparse tubules. **B)** Effect of altered surface/inner CRU NCX expression ratio on voltage (i) and global cytosolic Ca^2+^ concentration (ii) in cells with sparse (left), intermediate (middle), and dense (right) tubules following pacing at 3 Hz to examine the occurrence of DADs and SCRs. **C)** Mechanism underlying increasing surface/inner CRU NCX expression ratio inhibiting SCRs and DADs. Biomarkers were determined from the first 100 ms of no-stimulation period and normalized to those of cells with a retained surface/inner CRU NCX expression ratio of 1.0. Higher surface/inner CRU NCX expression ratio is associated with reduced NCX contribution to Ca^2+^ extrusion by inner coupled CRUs, resulting in higher cleft Ca^2+^ concentration in inner CRUs with no visible effect in surface CRUs. Higher inner cleft Ca^2+^ concentration in inner CRUs causes increased RyR P_o_ and RyR leak, leading to increased SCRs and DADs. Since fewer inner CRUs containing NCX exist in sparsely tubulated cells to begin with, the impact of varying NCX distribution is more pronounced in cells with tubules. As such, the consequence of changing cleft Ca^2+^ concentration is greater in cells with dense and intermediate tubules rather than those with sparse tubules, explaining the observed differences in SCRs and DADs between cells with varying tubule densities.

**Figure 5.**
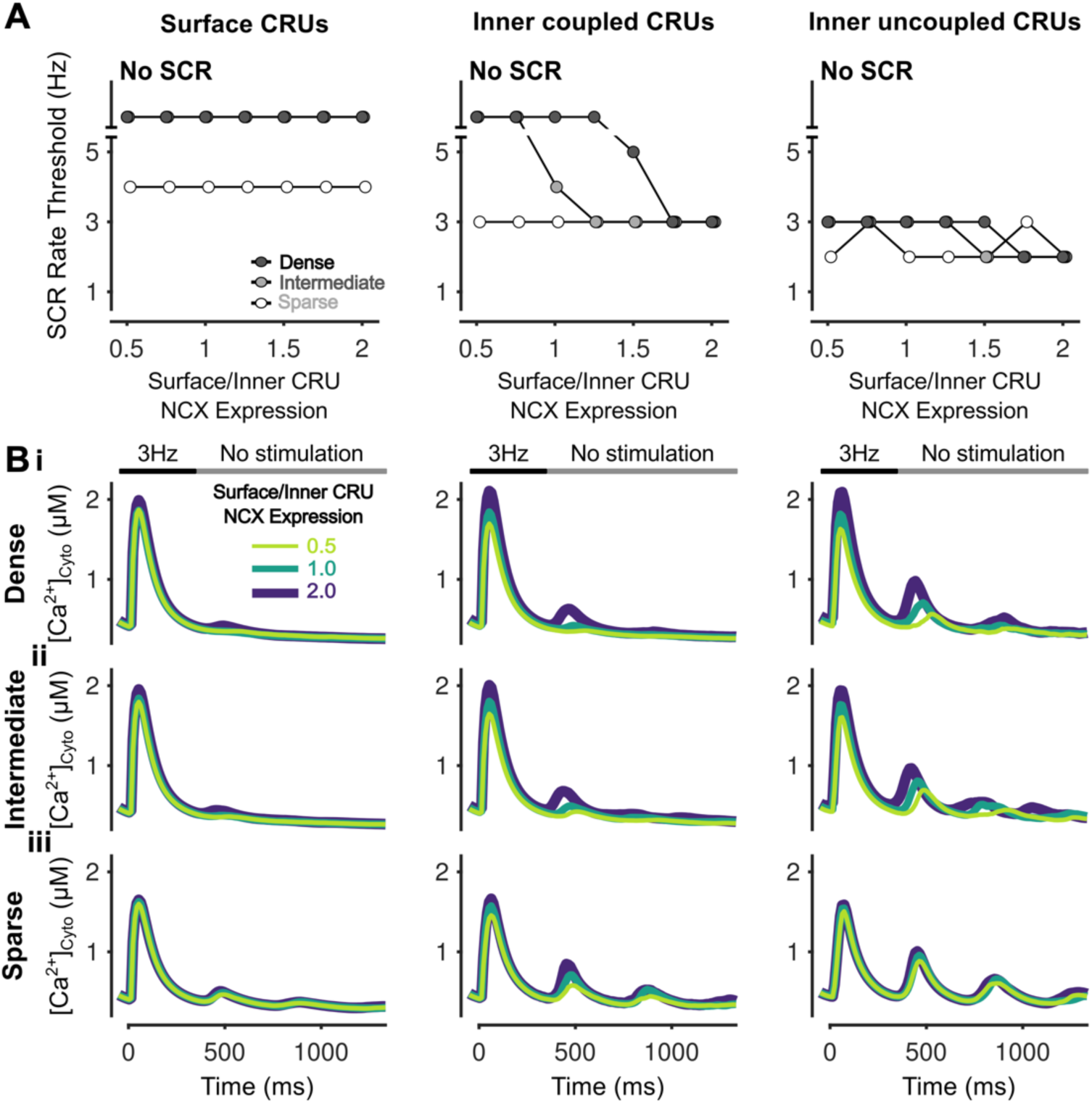
Increasing surface/inner CRU NCX expression ratio inhibits simultaneous SCRs in inner coupled CRUs. **A)** Rate threshold for local SCRs of surface (left), inner coupled (middle) and inner uncoupled (right) CRUs in sparse, intermediate and densely tubulated cells with the impact of altered surface/inner NCX expression ratio greatest in inner coupled CRUs of cells with dense and intermediate tubules. **B)** Cytosolic Ca^2+^ concentration of surface (left), inner coupled (middle) and inner uncoupled (right) CRUs from cells with dense (i), intermediate (ii), and sparse (iii) tubules following pacing at 3 Hz showing observed SCRs with varying surface/inner CRU NCX expression ratios.

### Inhibiting RyR has biphasic effects on SCRs, DADs, and SAPs, depending on the balance between the reduced number of RyRs and increased P_O_

To understand the effects of varying RyR expression and TATS density on SCRs, DADs, and SAPs, we altered RyR expression in cells with sparse, intermediate, and dense TATS as described in the Methods. Slight lowering of RyR expression above 70% of its baseline value was proarrhythmic and reduced the rate thresholds of SCRs, DADs, and SAPs. RyR expression below 40-70% typically had the opposite effect with rate threshold increased (**Fig. 6Ai-iii**). Interestingly, in the cell with sparse TATs the SCR rate threshold was unaffected by varying RyR expression levels, despite the ability of SCRs to generate DADs and SAPs being altered in these cells. Indeed, DADs and SAPs occur at overall lower pacing rates in cells with sparse TATS vs. those with intermediate or dense TATS, suggesting that TATS loss further promotes arrhythmogenic behavior (due to larger SCRs, especially in inner uncoupled CRUs, **Fig. 7B**). The biphasic effects of RyR inhibition on SCRs, DADs, and SAPs are shown in representative traces of voltage (**Fig. 6Bi**) and [Ca^2+^]_Cyto_ (**Fig. 6Bii**) following 4-Hz pacing. Our model analysis indicates that the biphasic effects of RyR inhibition are mediated by the balance between two contrasting processes: on one hand, reduction in RyR expression reduces SR Ca^2+^ release and increases the SR Ca^2+^ load (**Fig. 6Ci**), which elevates unitary RyR P_O_ (**Fig. 6Cii**); on the other hand, the reduced RyR number limits the magnitude of diastolic Ca^2+^ release by affecting the number of unitary release events and thus their ability to recruit neighboring RyRs and CRUs. As with the impact on global Ca^2+^, subcellular Ca^2+^ signaling also shows biphasic changes in SCR rate thresholds, which are most remarkable in surface CRUs of the cell with sparse TATS and in inner coupled CRUs of cells with intermediate and dense TATS (**Fig. 7A**). Interestingly, the rate threshold of inner coupled SCRs is similar to the whole cell SCR rate threshold (**Fig. 6Ai**), suggesting that SCRs in inner coupled CRUs dominantly affect whole cell SCR behavior. Furthermore, the rate threshold of surface SCRs (**Fig. 7A**) is similar to the rate thresholds of DADs and SAPs (**Fig. 6Aii-iii**), suggesting that surface CRU SCRs are more likely associated with the occurrence of DADs/SAPs.

**Figure 6.**
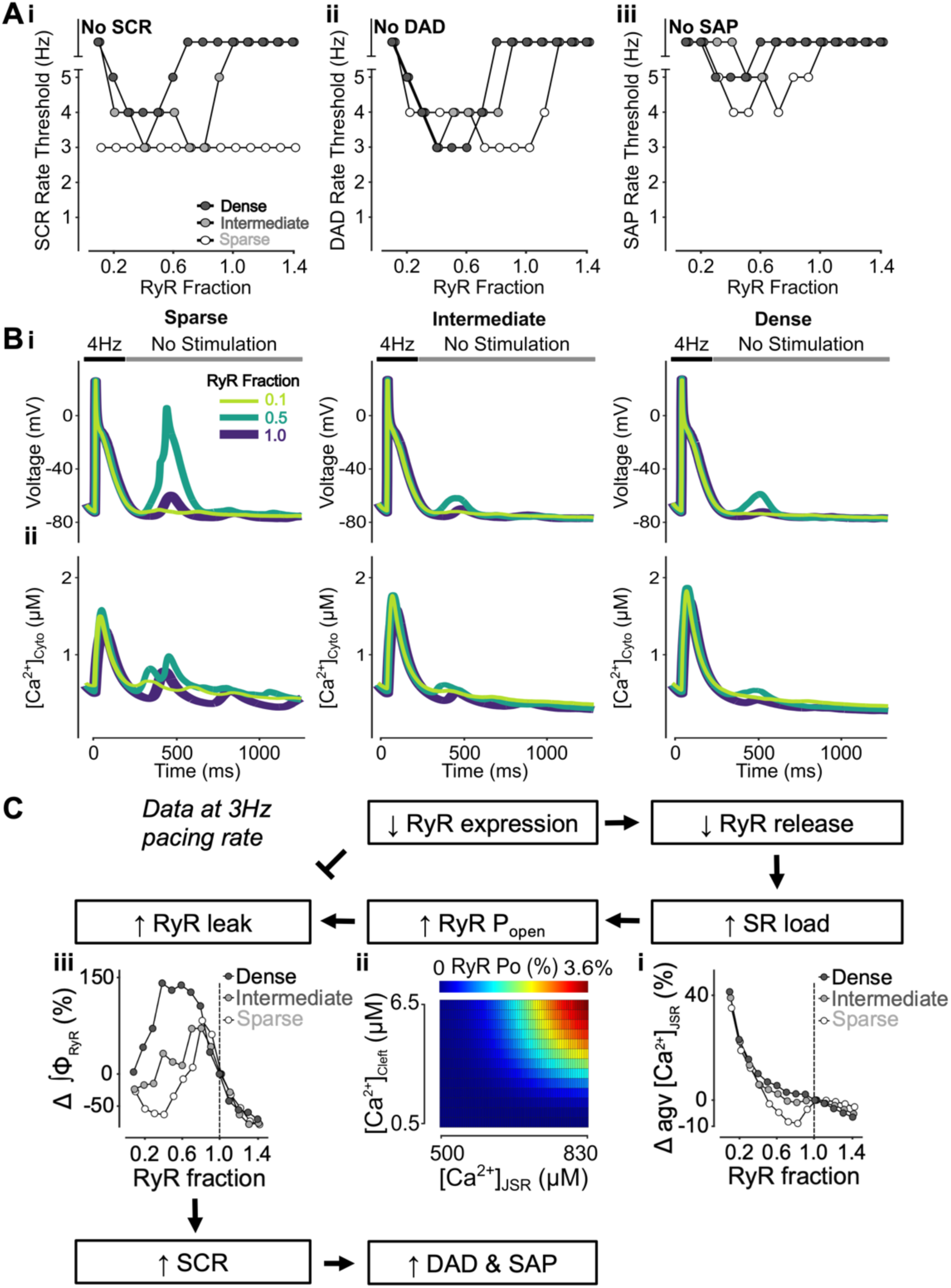
Inhibition of RyR has biphasic effects on SCRs, DADs, and SAPs. **A)** Rate threshold for SCRs (i), DADs (ii) and SAP (iii) in dense, intermediate and sparsely tubulated cells. Biphasic dependence on RyR inhibition is observed in all settings and cell types, with the exception of SCRs in sparsely tubulated cells. Here, the lowest observed baseline rate threshold for SCRs is unaltered by RyR inhibition. **B)** Effect of altered RyR fraction on voltage (i) and global cytosolic Ca^2+^ concentration (ii) in cells with sparse (left), intermediate (middle), and dense (right) tubules following pacing at 4 Hz to examine the occurrence of DADs, SAPs and SCRs. **C)** Mechanism underlying RyR inhibition having biphasic effects on SCRs, DADs and SAP. Biomarkers were determined from the first 100 ms of no-stimulation period and normalized to those of cells with a retained RyR fraction of 1.0. Lower RyR expression is associated with reduced RyR release, leading to elevated SR load (i) and augmented RyR P_o_ (ii). While higher RyR P_o_ causes RyR leak leading to increased SCRs, lower RyR expression itself inhibits RyR leak. This results in the biphasic effect of RyR reduction on SCRs, DADs and SAPs.

**Figure 7.**
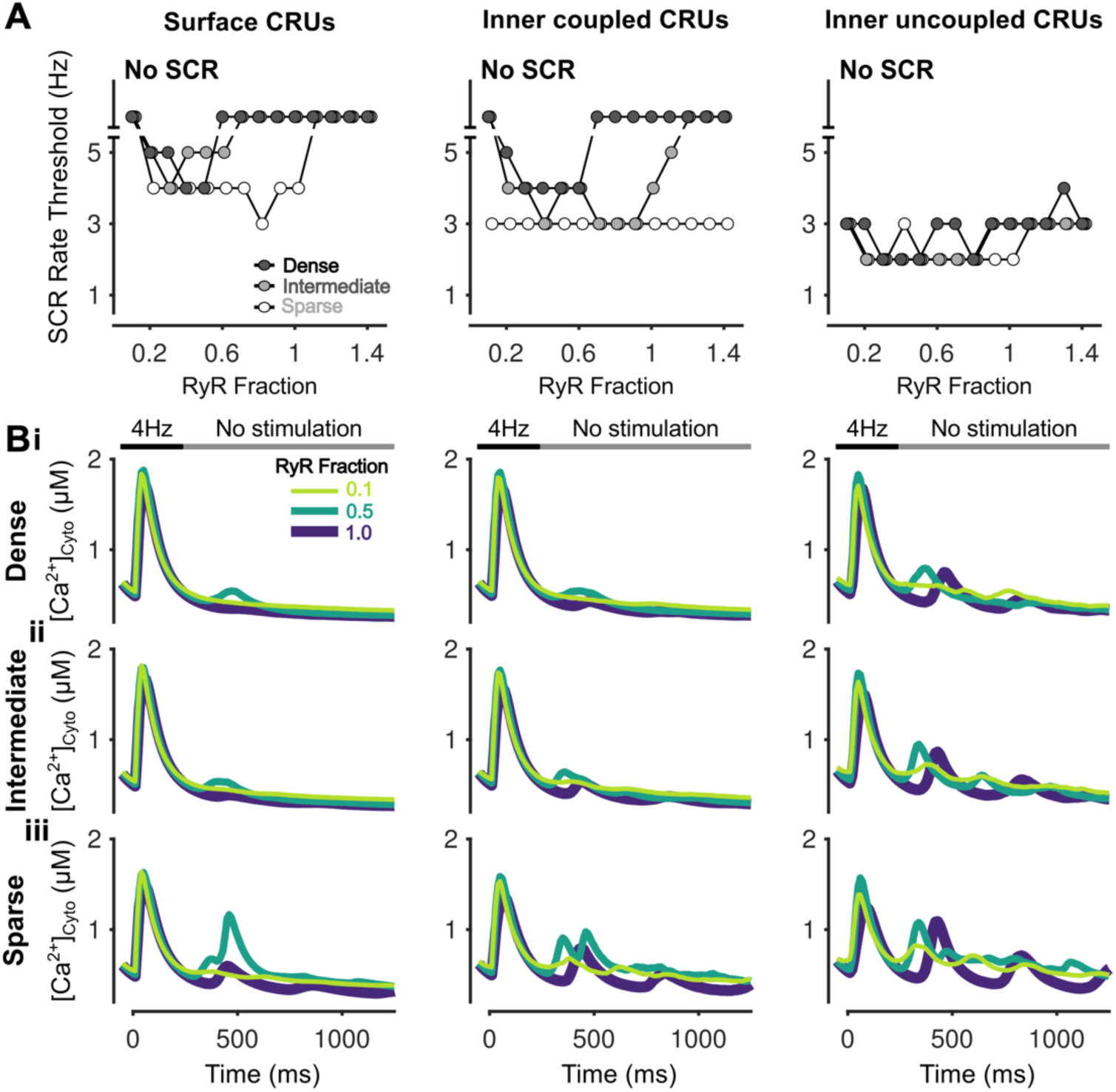
Inhibition of RyR has biphasic effects on SCRs in all CRUs. **A)** Rate threshold for local SCRs of surface (left), inner coupled (middle) and inner uncoupled (right) CRUs in sparse, intermediate and densely tubulated cells with the impact of altered RyR expression ratio greatest in surface CRUs of all cells and inner coupled CRUs of cells with dense and intermediate tubules. **B)** Cytosolic Ca^2+^ concentration of surface (left), inner coupled (middle) and inner uncoupled (right) CRUs from cells with dense (i), intermediate (ii), and sparse (iii) tubules following pacing at 4 Hz showing observed SCRs with varying RyR fraction.

### Increasing RyR expression in surface vs. inner CRUs inhibits SCRs but has modest effects on DADs and SAPs

To reveal the effects of heterogenous CRU RyR localization and TATS on SCRs, DADs, and SAPs, we varied the surface-to-inner CRU RyR expression ratio in cells with sparse, intermediate, and dense TATS. The simulation results indicate that increasing the surface/inner CRU RyR expression ratio inhibits SCRs in the cell with sparse and intermediate TATS, with no changes observed in the densely tubulated cell (**Fig. 8Ai**). However, modest changes were seen in the DAD threshold and only for the most extreme changes (e.g., ratios of 0.5 and 1.75 - 2.0, **Fig. 8Aii**), while no changes were detected in the SAP rate threshold (**Fig. 8Aiii**). Both increasing and decreasing the RyR expression in surface vs. inner CRUs diminished DADs (i.e., reduced amplitude and increased pacing threshold) in the cell with sparse TATS, but enhanced DADs in the cell with intermediate TATS, as demonstrated in the representative traces (**Fig. 8Bi**). The representative traces of [Ca^2+^]_Cyto_ also show that SCRs are larger in cells with sparse/intermediate vs. dense TATS (**Fig. 8Bii**) with varying surface/inner CRU RyR expression, as we show in the companion paper (Zhang et al.). The mechanistic analysis indicates increasing surface/inner CRU RyR expression enhances NCX-RyR coupling, thus leading to both larger NCX Ca^2+^ extrusion (**Fig. 8Ci**), with subsequent diminished SCRs that would attenuate DADs, and enhanced ΔV_m_/ΔCa^2+^ gain that would favor DADs (**Fig. 8Cii**). The balance of these two competing processes underlies the modest effects of varying RyR relative distribution in surface and inner CRUs on DADs. The analysis of subcellular Ca^2+^ signaling indicates that increasing surface RyR expression increases the SCR rate threshold in all CRUs, especially in inner CRUs, and that TATS loss exacerbates the effects of reducing RyR expression in inner CRUs (**Fig. 9A-B**).

**Figure 8.**
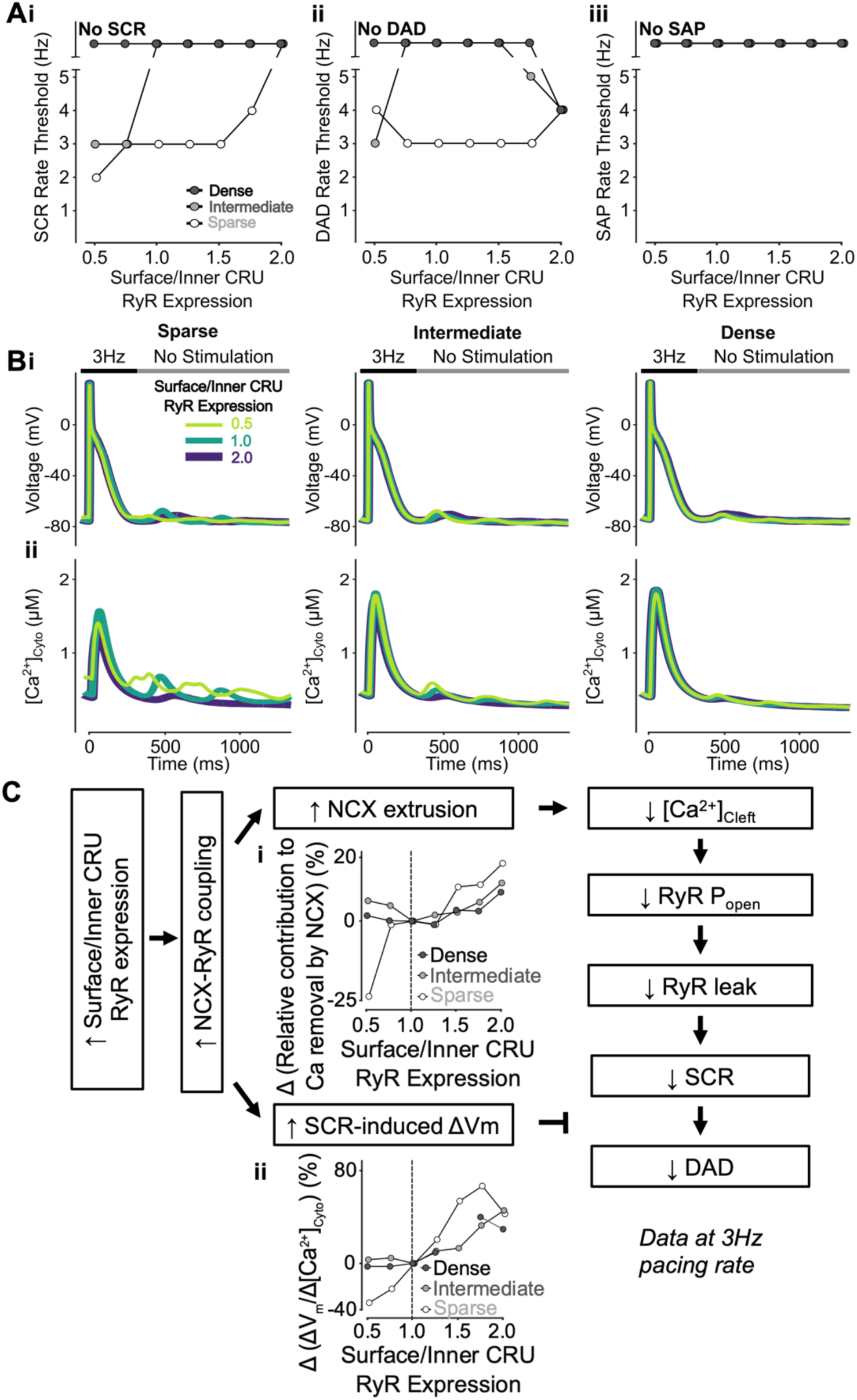
Increasing surface/inner CRU RyR expression ratio inhibits SCRs, has biphasic effects on DADs but does not affect SAP. **A)** Increasing surface/inner CRU RyR expression ratio (i.e., increasing RyR density in surface CRUs without changing whole-cell RyR expression) monotonically increases the rate threshold of SCRs (i), whereas DADs (ii) display biphasic dependence and SAP (iii) remains absent in all conditions. The effect on SCRs is greater in cells with sparse and intermediate tubular structures vs cells with dense tubules, whereas the biphasic effect on DADs is less pronounced in cells with sparse tubular structures vs cells with intermediate and dense tubules. **B)** Effect of altered surface/inner CRU RyR expression ratio on voltage (i) and global cytosolic Ca^2+^ concentration (ii) in cells with sparse (left), intermediate (middle), and dense (right) tubules following pacing at 3 Hz to examine the occurrence of DADs and SCRs. **C)** Mechanism underlying increasing surface/inner CRU RyR expression ratio inhibiting SCRs, having biphasic effects on DADs but not affecting SAP. Biomarkers were determined from the first 100 ms of no- stimulation period and normalized to those of cells with a retained surface/inner CRU RyR expression ratio of 1.0. Increasing surface/inner CRU RyR expression ratio means more RyRs are located closer to NCX, resulting in increased NCX-RyR coupling. This is associated with increased NCX contribution to Ca^2+^ extrusion (i) and higher SCR-induced voltage changes/SCR amplitude ratio (ii). Enhanced NCX extrusion results in lower cleft Ca^2+^ concentration, smaller RyR P_o_ and leak leading to milder SCRs. While this in itself limits DADs, DAD likelihood is promoted by the increased SCR-induced changes in V_m_, thus explaining the biphasic effects of increasing surface/inner CRU RyR expression ratio.

**Figure 9.**
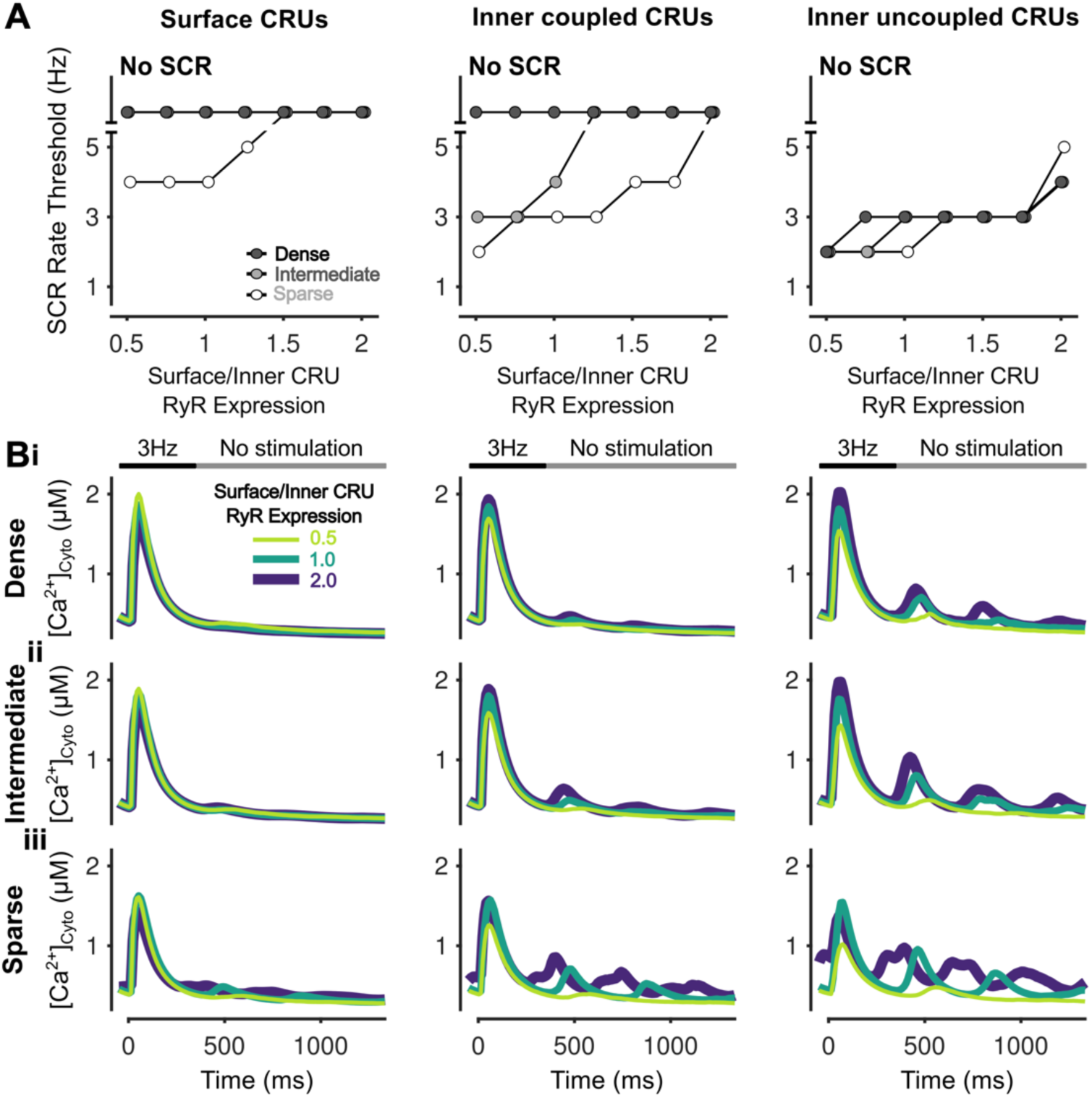
Increasing surface/inner CRU RyR expression ratio inhibits simultaneous SCRs in all CRUs. **A)** Rate threshold for local SCRs of surface (left), inner coupled (middle) and inner uncoupled (right) CRUs in sparse, intermediate and densely tubulated cells with the impact of altered surface/inner RyR expression ratio greatest in inner coupled CRUs of cells with sparse and intermediate tubules. **B)** Cytosolic Ca^2+^ concentration of surface (left), inner coupled (middle) and inner uncoupled (right) CRUs from cells with dense (i), intermediate (ii), and sparse (iii) tubules following pacing at 3 Hz showing observed SCRs with varying surface/inner CRU RyR expression ratios.

### Loss of CSQ promotes SCRs and DADs, primarily through diminished CSQ Ca^2+^ buffering, with no effect on SAPs

The impact of CSQ expression levels and TATS density on SCRs, DADs, and SAPs was examined by varying the expression of CSQ in cells with sparse, intermediate, and dense TATS. Because CSQ both affects luminal Ca^2+^ buffering and RyR P_O_, we simulated the changes in CSQ expression level by 1) only changing the Ca^2+^ buffering parameters (**Figs. 10-11**), 2) only modifying the parameters associated with CSQ regulation of RyR gating (**Figs. 11-12**), and 3) modulating both processes (**Figs. 13-14**), as described in the Methods. Reducing CSQ-mediated luminal SR Ca^2+^ buffering promotes SCRs, DADs, and SAPs (**Figs. 10A-B**), and enhances subcellular SCRs in all CRUs (**Fig. 11**). However, inhibiting CSQ RyR-regulation, i.e., decreasing SR Ca^2+^ sensitivity of RyR P_O_, conversely increases pacing rate thresholds of SCRs, DADs, and SAPs, especially in cells with sparse TATS (**Fig. 12A-B**). Inhibiting CSQ RyR-regulation also reduces subcellular SCRs in all CRUs, especially in inner CRUs (**Fig. 13A-B**). When concomitantly simulating the changes in both luminal Ca^2+^ buffering and RyR regulation, we found that CSQ inhibition promotes SCRs and DADs, without changing the SAP rate threshold (**Fig. 14A-B**), i.e., similar to the effects of inhibiting CSQ Ca^2+^ buffering alone (**Fig. 10A-B**). Indeed, our mechanistic analysis suggests that luminal Ca^2+^ buffering by CSQ plays a more important role in affecting arrhythmic outcomes than its role on RyR gating. On one hand, inhibiting CSQ Ca^2+^ buffering (**Fig. 14C, 1**) increased the SR load (**Fig. 14Ci**) and RyR leak (**Fig. 14Cii**) to promote SCRs and DADs. On the other hand, inhibition of CSQ RyR-regulation (**Fig. 14C, 2**) elevated RyR systolic P_O_ and enhanced SR release fraction, and decreased SR load (**Fig. 14Ciii**), leading to subsequent smaller RyR leak (**Fig. 14Civ**) and reduced SCRs and DADs, especially in inner coupled CRUs (**Fig. 15**). When combining the two opposing effects, we found an overall net increase in RyR leak (**Fig. 14Cvi**), which was dominant over the slight decrease in SR load (**Fig. 14Cv**), and promoted SCRs, DADs, and SAPs. In these settings, the rate thresholds of SCRs, DADs, and SAPs are consistently lower in the cell with sparse TATS (in agreement with results in the companion Zhang et al. paper), but cells with intermediate and dense TATS are more susceptible to changes in CSQ expression.

**Figure 10.**
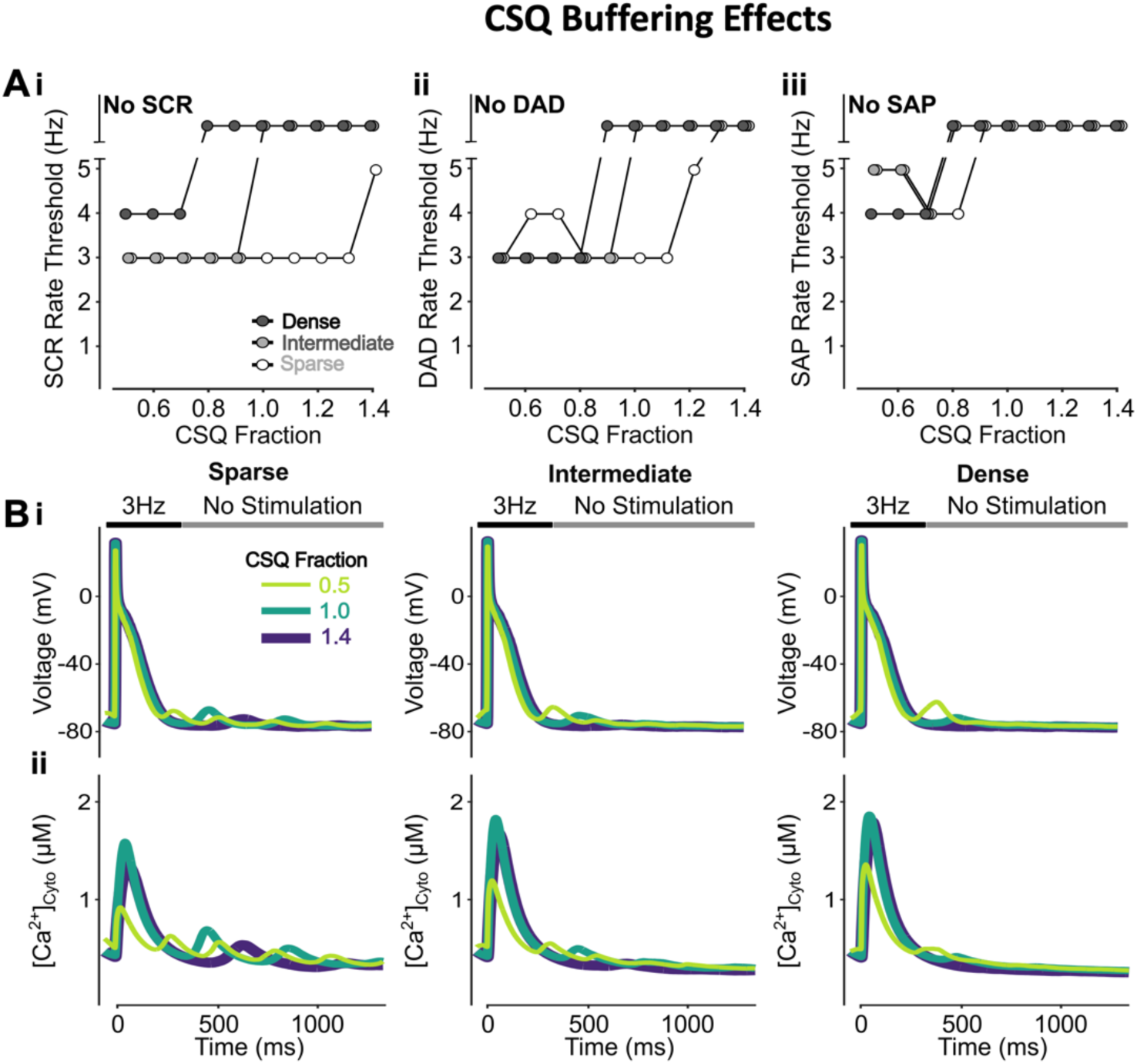
Promotion of CSQ Ca^2+^ buffering effects inhibits SCRs, DADs and SAPs. **A)** Promoting Ca^2+^ buffering by CSQ monotonically increases the rate threshold of SCRs (i), DADs (ii), and SAPs (iii). This effect is enhanced in cells with dense and intermediate tubular structures vs cells with sparse tubules. **B)** Effect of altered CSQ-mediated Ca^2+^ buffering on voltage (i) and global cytosolic Ca^2+^ concentration (ii) in cells with sparse (left), intermediate (middle), and dense (right) tubules following pacing at 3 Hz to examine the occurrence of DADs and SCRs.

**Figure 11.**
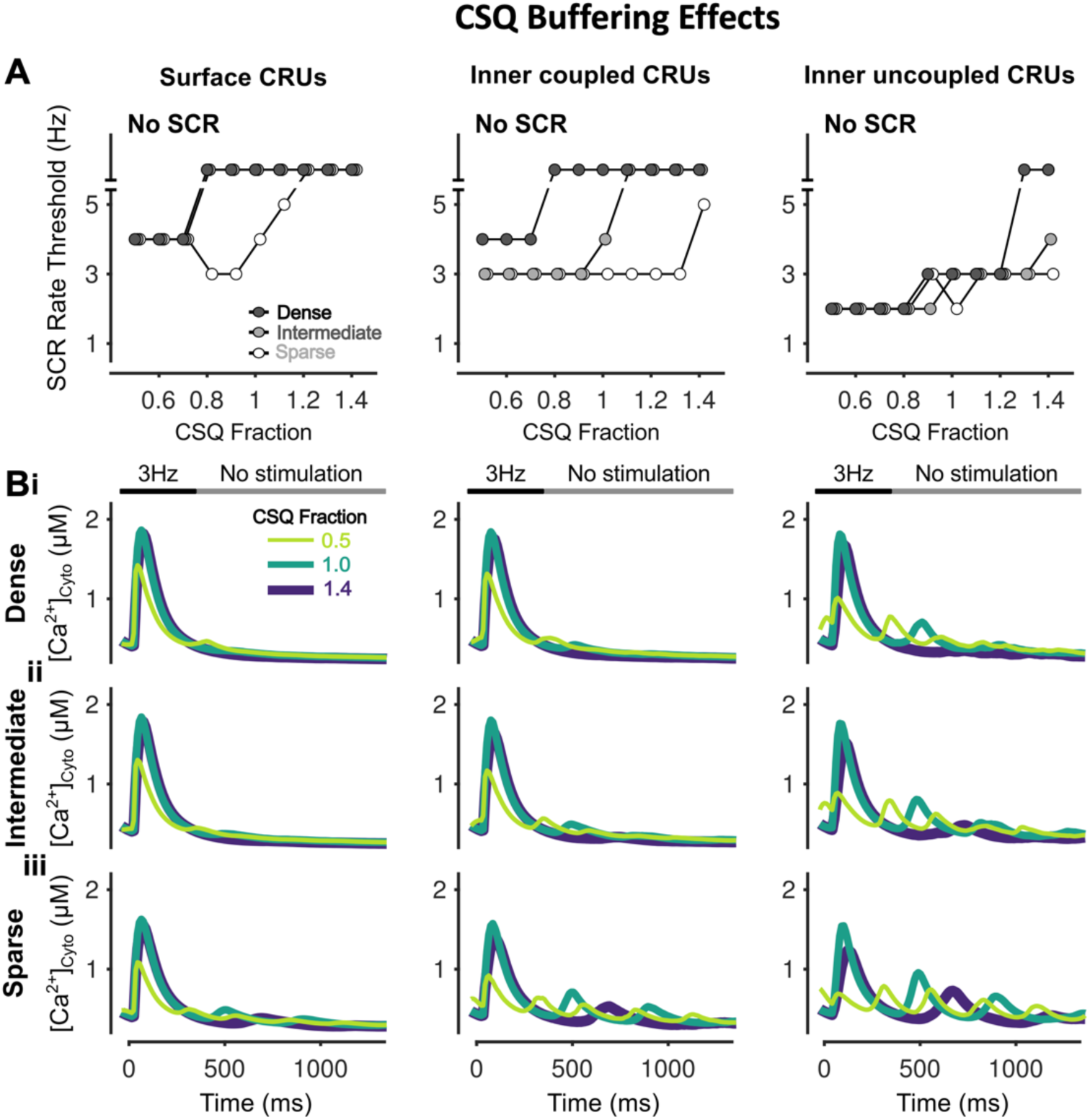
Inhibition of CSQ Ca^2+^ buffering effects promotes SCRs in all CRUs. **A)** Rate threshold for local SCRs of surface (left), inner coupled (middle) and inner uncoupled (right) CRUs in sparse, intermediate and densely tubulated cells with the impact of altered CSQ Ca^2+^ buffering effects greatest in inner coupled/uncoupled CRUs of cells with dense and intermediate tubules. **B)** Cytosolic Ca^2+^ concentration of surface (left), inner coupled (middle) and inner uncoupled (right) CRUs from cells with dense (i), intermediate (ii), and sparse (iii) tubules following pacing at 3 Hz showing observed SCRs with varying fraction of CSQ Ca^2+^ buffering effects.

**Figure 12.**
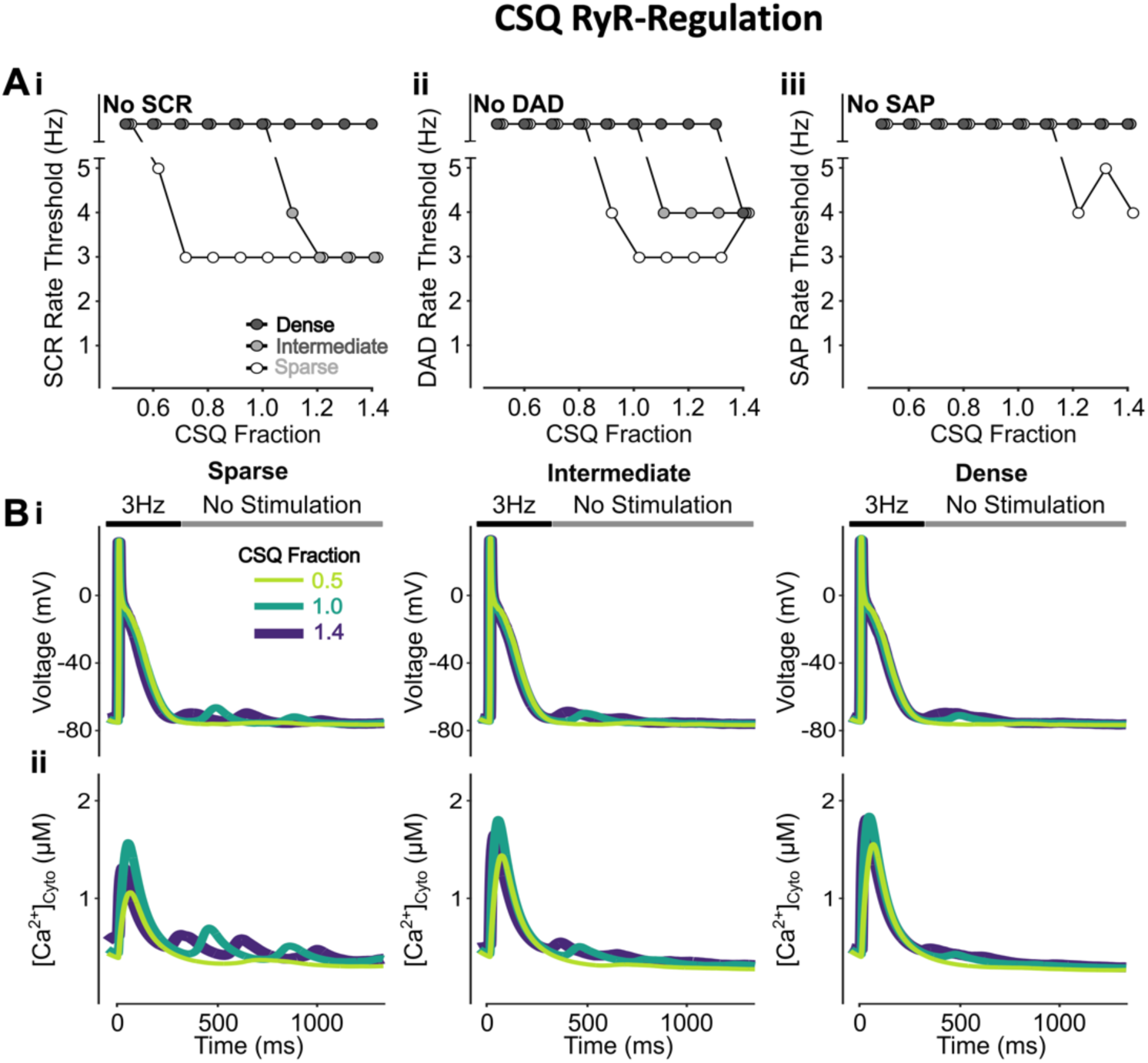
Increasing CSQ-RyR regulation promotes SCRs, DADs and SAPs. **A)** Increasing CSQ-RyR regulation monotonically decreases the rate threshold of SCRs (i), DADs (ii), and SAPs (iii) with the effect greater in cells with sparse and intermediate tubular structures vs cells with dense tubules. **B)** Effect of altered CSQ-RyR regulation on voltage (i) and global cytosolic Ca^2+^ concentration (ii) in cells with sparse (left), intermediate (middle), and dense (right) tubules following pacing at 3 Hz to examine the occurrence of DADs and SCRs.

**Figure 13.**
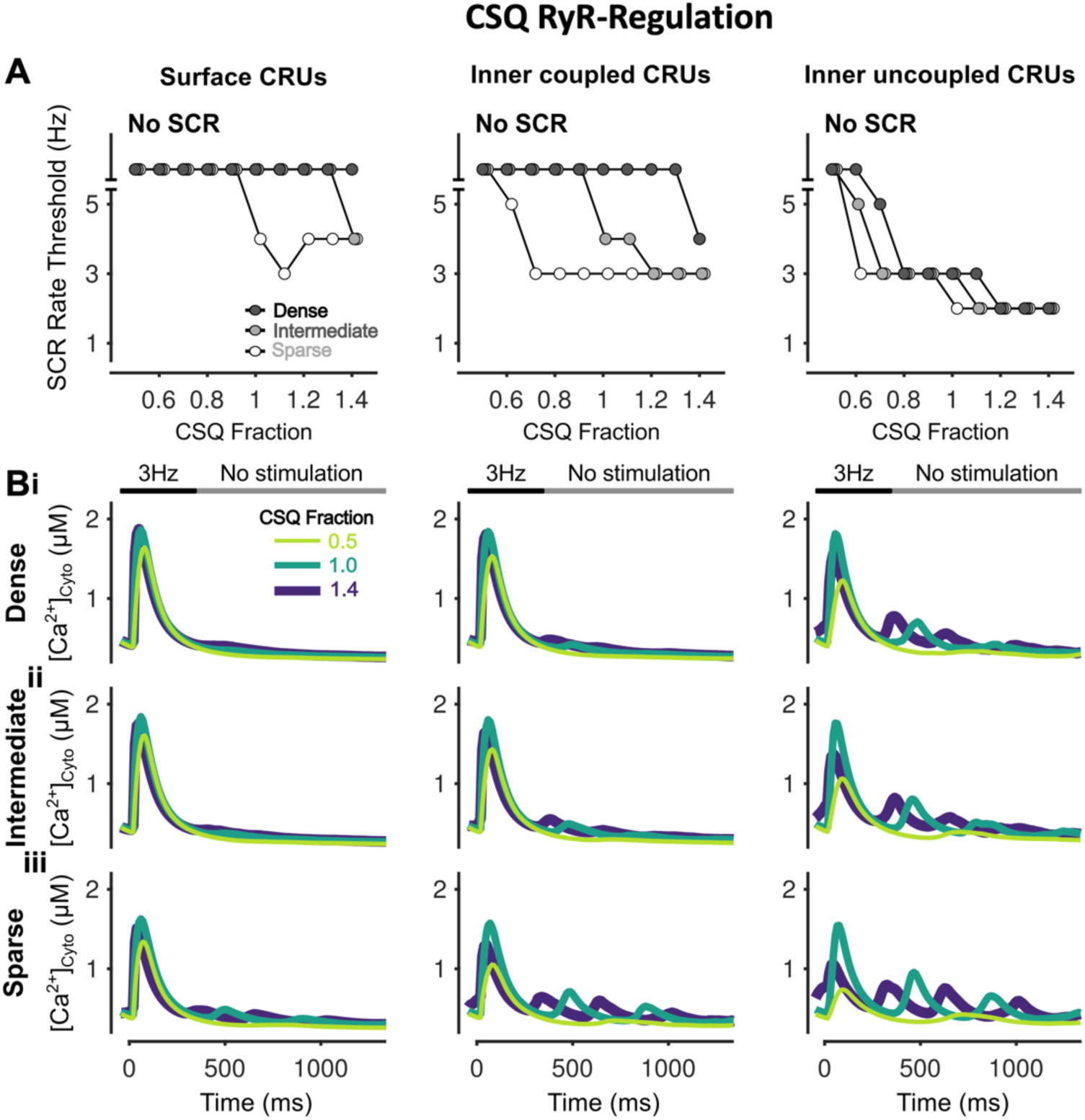
Inhibition of CSQ-RyR regulation suppresses SCRs in all CRUs. **A)** Rate threshold for local SCRs of surface (left), inner coupled (middle) and inner uncoupled (right) CRUs in sparse, intermediate and densely tubulated cells with the impact of altered CSQ- RyR regulation greatest in inner coupled/uncoupled CRUs. **B)** Cytosolic Ca^2+^ concentration of surface (left), inner coupled (middle) and inner uncoupled (right) CRUs from cells with dense (i), intermediate (ii), and sparse (iii) tubules following pacing at 3 Hz showing observed SCRs with varying fraction of CSQ RyR-regulation.

**Figure 14.**
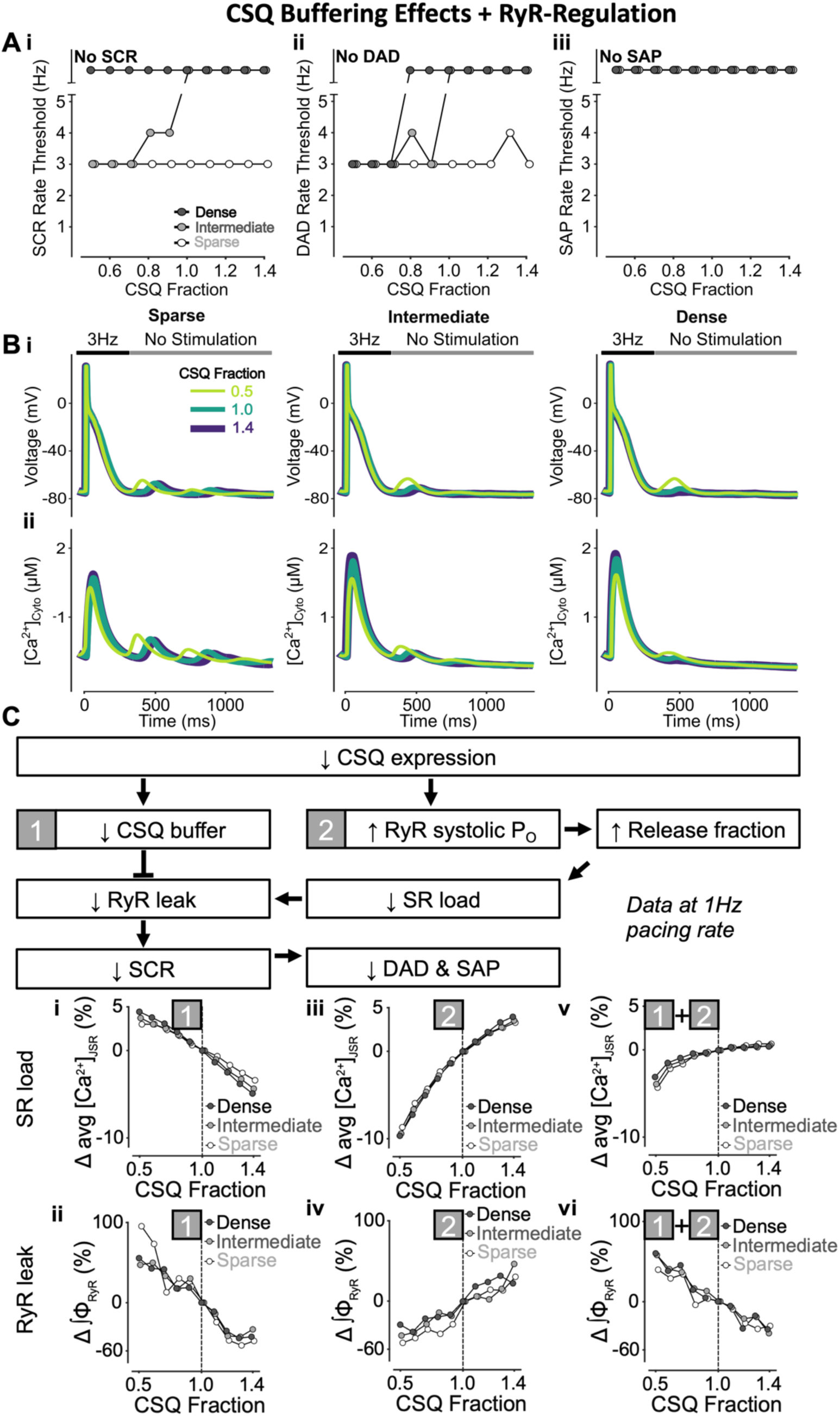
Increasing CSQ fraction inhibits SCRs and DADs with no effect on SAPs. **A)** Increasing CSQ fraction (i.e., both CSQ-mediated Ca^2+^ buffering and RyR regulation) monotonically raises the rate threshold of SCRs (i) and DADs (ii), with SAP (iii) remaining absent in all conditions. While CSQ-mediated changes in SCRs are solely observed in cells with intermediate tubular structures, the effect of reduced CSQ fraction on DADs is greater in cells with dense and intermediate tubular structures vs cells with sparse tubules. **B)** Effect of altered CSQ fraction on voltage (i) and global cytosolic Ca^2+^ concentration (ii) in cells with sparse (left), intermediate (middle), and dense (right) tubules following pacing at 3 Hz to examine the occurrence of DADs and SCRs. **C)** Mechanism underlying increased CSQ fraction inhibiting SCRs and DADs without affecting SAPs. Biomarkers were determined from the first 100 ms of no-stimulation period and normalized to those of cells with a retained CSQ fraction of 1.0. Since CSQ is a jSR Ca^2+^ buffer and regulates RyR P_o_, there is interplay between varying CSQ buffering effects (1), CSQ-RyR regulation (2) and CSQ fraction (1+2). Lower CSQ expression is associated with reduced CSQ-mediated jSR Ca^2+^ buffering (1) and increased RyR systolic P_o_ (2). Reduced CSQ Ca^2+^ buffering (1) decreases SR load (i) but promotes RyR leak (ii), and thus SCRs. Conversely, enhanced systolic RyR P_o_ (2) increases SR release fraction, which lowers SR load (iii) and diminishes RyR leak (iv), leading to reduced SCRs. Combining these two effects, although decreasing CSQ expression lowers down SR Ca^2+^ load (v), RyR leak is enhanced (vi), which leads to promotion of SCRs, DADs, and SAPs.

**Figure 15.**
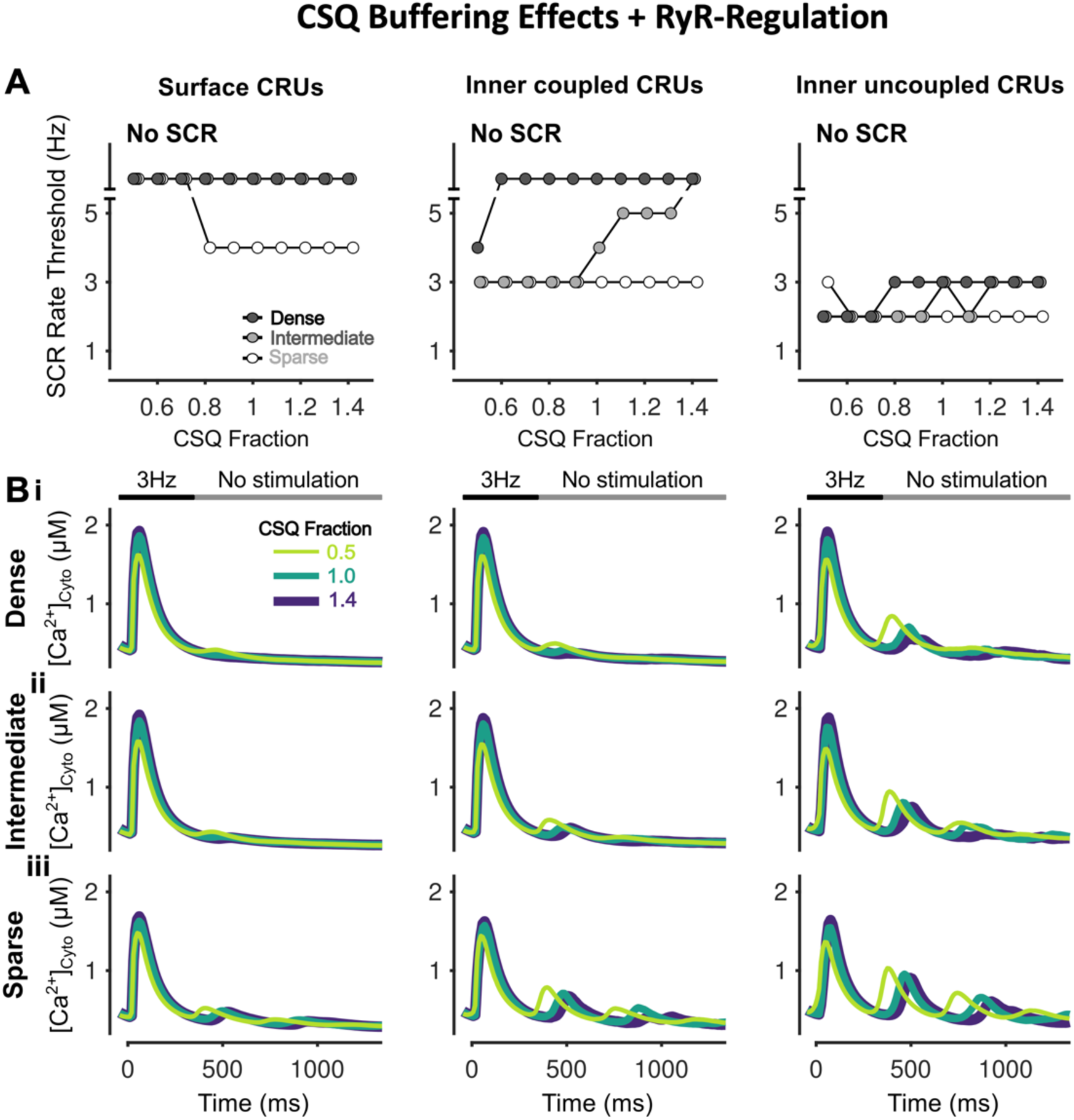
Varying CSQ fraction has differing effects on SCR dependent on CRU location. **A)** Rate threshold for local SCRs of surface (left), inner coupled (middle) and inner uncoupled (right) CRUs in sparse, intermediate and densely tubulated cells. Whereas CSQ downregulation raises the SCR rate threshold in cells with sparse tubules, it lowers the threshold in inner coupled/uncoupled CRUs of cells with intermediate and dense tubular structures. **B)** Cytosolic Ca^2+^ concentration of surface (left), inner coupled (middle) and inner uncoupled (right) CRUs from cells with dense (i), intermediate (ii), and sparse (iii) tubules following pacing at 3 Hz showing observed SCRs with varying CSQ fraction.

### Increasing surface vs. inner CRU CSQ expression promotes SCRs but inhibits DADs by elevating inner RyR leak but decreasing surface RyR leak

To reveal the effects of heterogeneous subcellular CSQ distribution and TATS density on SCRs, DADs, and SAPs, we simulated various surface/inner CRU CSQ expression ratios in cells with sparse, intermediate, and dense TATS. The simulation results indicate that increasing surface vs. inner CRU CSQ expression lowers the pacing threshold for SCRs (in cells with intermediate TATS, **Fig. 16Ai**), but increased the pacing threshold for DADs (in cells with sparse and intermediate TATS, **Fig. 16Aii**), without remarkable effects on SAP rate threshold (**Fig. 16Aiii**). These results are also illustrated by the representative traces of [Ca^2+^]_Cyto_ and V_m_ (**Fig. 16B**). We found that increasing surface-to-inner CRU CSQ expression ratio changes RyR leak locally, i.e., decreasing RyR leak at the periphery and increasing it at the cell interior. A larger inner RyR leak favors SCRs and DADs (**Fig. 16Ci**), but a smaller RyR leak in the periphery area (**Fig. 16Cii**) reduces NCX Ca^2+^ extrusion (**Fig. 16Ciii**), ΔV_m_/ΔCa^2+^ gain (**Fig. 16Civ**) and DADs. The effects of varying surface-to-inner CRU CSQ expression ratio on ΔV_m_/ΔCa^2+^ gain are stronger in cells with sparse TATS (**Fig. 16Civ**), which is reflected in DAD rate threshold changes being more marked compared to intermediate/densely tubulated cells (**Fig. 16Aii**). Examination of the subcellular Ca^2+^ signals suggests that increasing surface vs. inner CSQ expression reduces the SCR rate threshold in inner coupled CRUs of the cell with intermediate TATS (**Fig. 17A**), matching the changes in SCR rate threshold (**Fig. 16Ai**), and in all cases, SCRs are favored in cells with sparse TATS (**Fig. 17A-B**).

**Figure 16.**
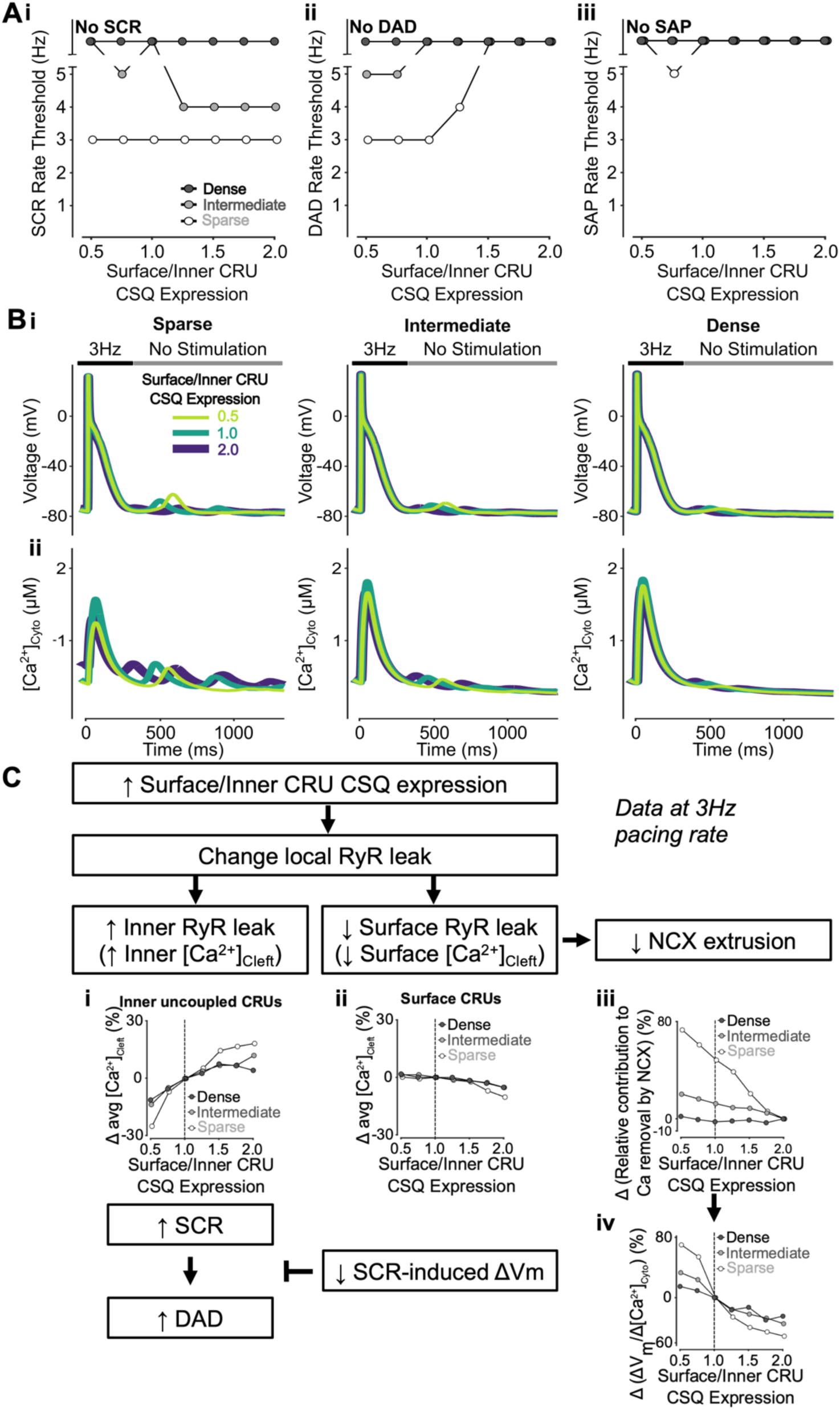
Increasing surface/inner CRU CSQ expression ratio promotes SCRs but inhibits DADs and does not affect SAP. **A)** Increasing surface/inner CRU CSQ expression ratio (i.e., increasing CSQ density in surface CRUs without changing whole-cell CSQ expression) monotonically decreases the rate threshold of SCRs (i) and increases the rate threshold of DADs (ii), whereas SAPs (iii) only appear in cells with sparse tubules and surface/inner CRU CSQ expression ratio of 0.75. The effect on SCRs is greater in cells with intermediate tubular structures vs cells with sparse and dense tubules, whereas the effect on DADs is enhanced in cells with sparse and intermediate tubular structures vs cells with dense tubules. **B)** Effect of altered surface/inner CRU CSQ expression ratio on voltage (i) and global cytosolic Ca^2+^ concentration (ii) in cells with sparse (left), intermediate (middle), and dense (right) tubules following pacing at 3 Hz to examine the occurrence of DADs and SCRs. **C)** Mechanism underlying increasing surface/inner CRU CSQ expression ratio promoting SCRs, inhibiting DADs with no effect on SAPs. Biomarkers were determined from the first 100 ms of no-stimulation period and normalized to those of cells with a retained surface/inner CRU CSQ expression ratio of 1.0. As similarly described when altering overall CSQ expression (Figure 13), increasing surface/inner CRU CSQ expression ratio is associated with increased RyR leak in inner uncoupled CRUs (i) and decreased RyR leak in surface CRUs (ii). Enhanced RyR leak in inner uncoupled CRUs leads to stronger SCRs and DADs. However, less RyR leak in surface CRUs reduces NCX contribution to Ca^2+^ extrusion (iii) and lowers SCR- induced voltage changes/SCR amplitude ratio (iv). As such, while SCRs typically lead to DADs, this transition is prohibited by the decreased SCR-induced changes in V_m_, thus explaining the opposing effects of increasing surface/inner CRU RyR expression ratio on SCRs and DADs.

**Figure 17.**
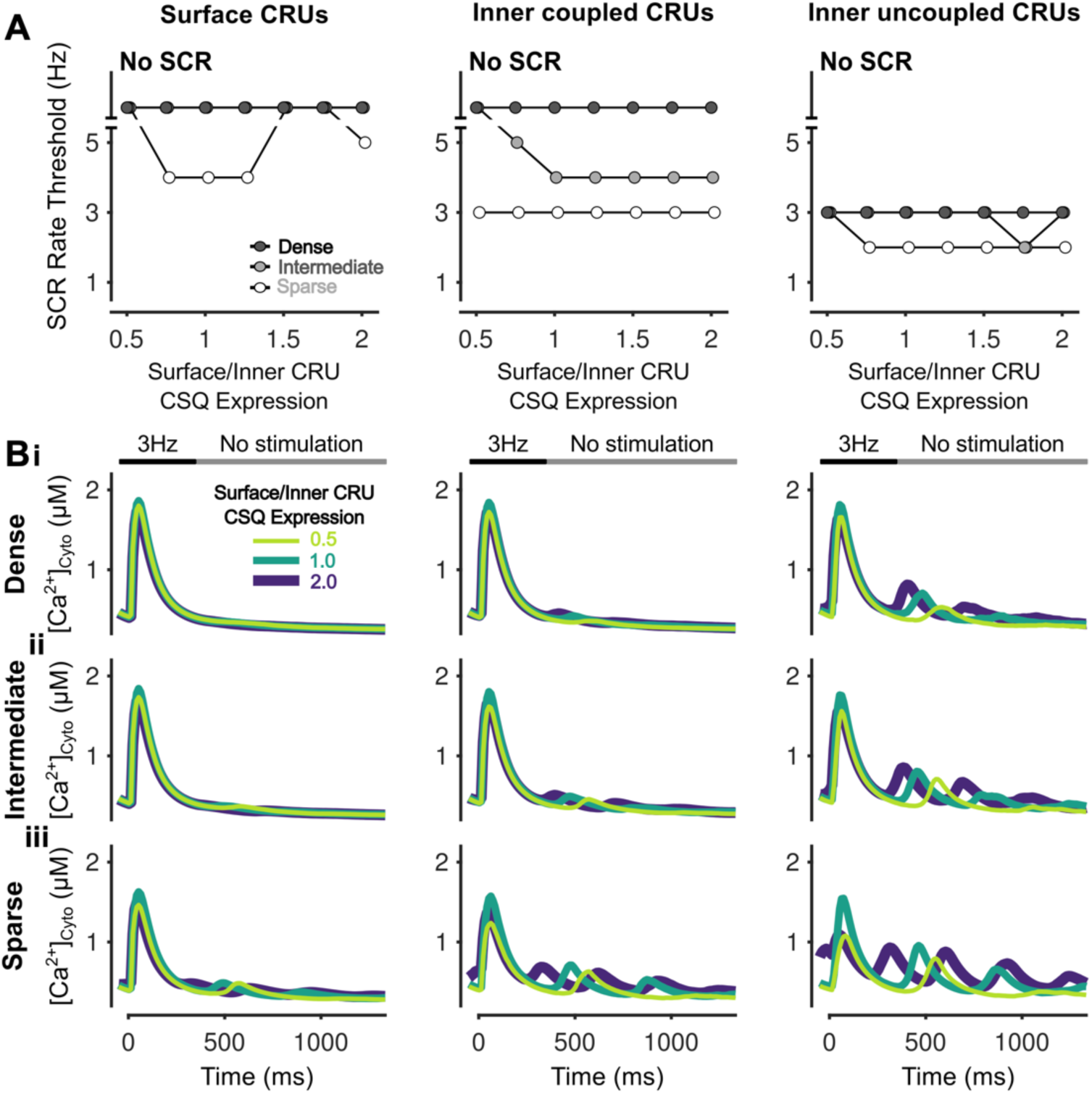
Increasing surface/inner CRU CSQ expression ratio promotes simultaneous SCRs in inner coupled CRUs of cells with intermediate tubules. **A)** Rate threshold for local SCRs of surface (left), inner coupled (middle) and inner uncoupled (right) CRUs in sparse, intermediate and densely tubulated cells with the impact of altered surface/inner CSQ expression ratio greatest in inner coupled CRUs of cells with intermediate tubules. B) Cytosolic Ca^2+^ concentration of surface (left), inner coupled (middle) and inner uncoupled (right) CRUs from cells with dense (i), intermediate (ii), and sparse (iii) tubules following pacing at 3 Hz showing observed SCRs with varying surface/inner CRU CSQ expression ratios.

## Discussion

In this study, we utilized our integrative model of the human atrial myocyte (Zhang et al.) to predict and quantitatively explain how TATS loss interacts with varying expression and localization of key Ca^2+^ handling proteins to disrupt diastolic Ca^2+^ homeostasis and electrophysiological stability in human atrial cells. Specifically, we demonstrated that arrhythmogenic effects of Ca^2+^ handling protein remodeling are especially exaggerated in cells with intermediate TATS, with densely tubulated myocytes being more resilient, and detubulated myocytes being rather insensitive to the superimposed ionic remodeling. These findings provide mechanistic insight into the interactive contributions of TATS and Ca^2+^-handling protein expression and distribution on Ca^2+^-driven proarrhythmic behavior that underlie AF pathophysiology and may help to predict the effects of antiarrhythmic strategies at varying stages of ultrastructural remodeling.

### NCX expression and distribution

In the healthy heart, NCX inhibition prevents Ca^2+^ extrusion, leading to Ca^2+^ overload (Bers, 2002) and thus enhancement of SCRs (Lotteau *et al*., 2021). This is seen in our simulations, whereby a decrease in NCX is associated with a reduction in the pacing threshold for SCRs. While increased SCRs are typically expected to promote V_m_ instabilities, drug-induced NCX block enhanced SCRs but reduced DADs in rabbit atrial myocyte experiments (Hohendanner *et al*., 2015). We also observed this in our simulations where inhibiting NCX (both globally or locally in the cell interior vs. periphery) varied the balance between increased SCRs and reduced ΔV_m_/ΔCa^2+^ gain that determines the net impact on SCRs, DADs, and SAPs (**Fig. 2**). As such, our model provides quantitative spatially-detailed insight into the interaction of these two opposing effects. Furthermore, our simulation results suggest that SCR is further enhanced with TATS loss, and the effects of inhibiting NCX on [Ca^2+^]_Cyto_-voltage coupling is stronger in cells with intermediate and dense TATS (**Figs. 2** and **4**). This suggests that with severe ultrastructural remodeling and TATS loss, additional changes in Ca^2+^ extrusion may not strongly modulate Ca^2+^ homeostasis and arrhythmogenesis. Interestingly, atrial myocytes from NCX knock-out mice also have reduced TATS density (Yue *et al*., 2017). It is reasonable to speculate that TATS loss in this setting may be a compensatory adaptation to Ca^2+^ overload, to limit Ca^2+^-V_m_ coupling and attenuate DAD and SAP risk. Indeed, this is supported by our sparsely tubulated myocyte model having a higher SAP rate threshold than the models with intermediate and dense TATS.

Interestingly, lower NCX promotes CaT alternans in cells with sparse and intermediate TATS but has biphasic effects on the rate threshold of CaT alternans in the densely tubulated cell (**Fig. 18Ai**), with the same effect observed in all CRU locations (**Fig. 3Bi**). Similarly, NCX inhibition has biphasic effects on the rate threshold of APD alternans (**Fig. 18Aii**), alongside changes in rate thresholds of DADs and SAPs (**Fig. 2Aii-iii**). Though these effects were observed when altering NCX expression, varying surface/inner CRU NCX localization had no effect on rate threshold of APD alternans besides initiating some CaT alternans at 5Hz (**Fig. 18B**). Taken together, these results suggest that atrial cells are more sensitive to whole-cell rather than subcellular changes in NCX-mediated Ca^2+^ extrusion.

**Figure 18.**
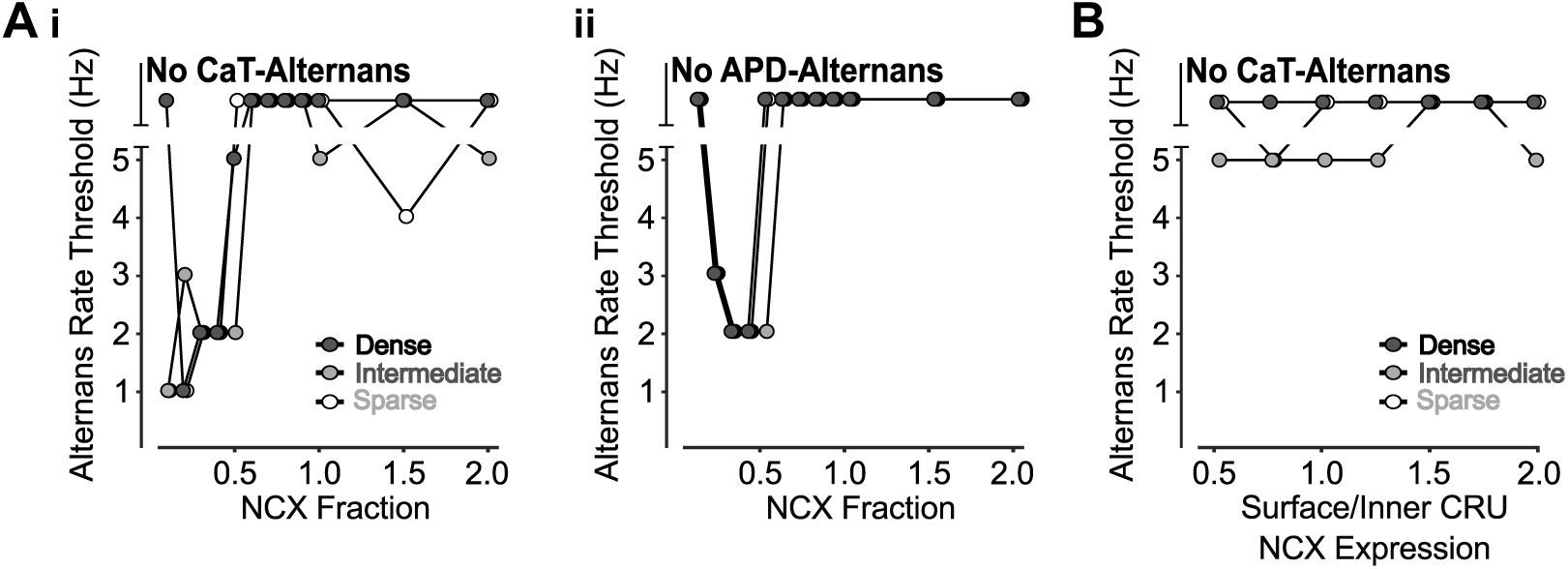
Inhibition of NCX has biphasic effects on CaT and APD alternans but increasing surface/inner CRU NCX expression ratio only exhibits CaT alternans at 5 Hz. **A)** The rate thresholds of CaT alternans (i) and APD alternans (ii) display biphasic dependence on NCX reduction. B) Varying surface/inner CRU NCX expression ratio induces CaT alternans at 5 Hz, especially in cells with intermediate TATS.

As NCX is upregulated in disease (Voigt *et al*., 2012), we also considered the impact of increased, not just decreased, NCX expression on arrhythmogenesis. Based on our simulation results, enhanced NCX expression may be an adaptive ionic remodeling process to protect the cells against Ca^2+^ overload-induced arrhythmia, although this can also be proarrhythmic with increased Ca^2+^-V_m_ coupling. Indeed, in AF, enhanced NCX expression increases diastolic Ca^2+^- V_m_ coupling to augment the arrhythmogenic effects of enhanced SR Ca^2+^-leak, causing transient inward current and subsequent DADs (Hove-Madsen *et al*., 2004; Lenaerts *et al*., 2009; Neef *et al*., 2010; Voigt *et al*., 2012). Because of this, NCX inhibition is thought to be a suitable therapeutic AF treatment (Hobai & O’Rourke, 2004). Our simulation suggests that varying degrees of NCX inhibition could have opposite effects on arrhythmias with NCX inhibition treatment potentially having better effects in cells with intermediate and dense TATS than in cells with sparse TATS. In atrial myocytes, NCX likely distributes on both the cell surface and TATS (Melnyk *et al*., 2005; Scriven *et al*., 2010), with localization disrupted with TATS remodeling in disease. As such, altered NCX distribution alongside TATS loss promotes arrhythmia, while increasing NCX density within remaining TATS may preserve NCX-mediated Ca^2+^-extrusion and rescue the cells from Ca^2+^ overload.

### RyR expression and distribution

Previous computational studies using a ventricular myocyte model show that enhanced RyR expression *per se* increased SR Ca^2+^ leak, while the subsequent reduction SR Ca^2+^ load reduced RyR P_O_ (Sato *et al*., 2016), similar to experimental data showing that SR Ca^2+^ leak reduces SR load (Bers, 2002), which can reduce the RyR P_O_ (Sitsapesan & Williams, 1994). On the other hand, a recent experimental study found RyR loss-of-function mutations, which are associated with arrhythmia and sudden cardiac death, cause elevated SR Ca^2+^ load (Li *et al*., 2021), which can increase the RyR P_O_. These data suggest that increasing RyR expression might be pro- or anti-arrhythmic depending on the prevailing mechanism. Indeed, our simulations indicate that altering RyR expression shifts the balance between RyR number-associated release flux and opposite SR load changes that determine the net impact on RyR leak, and subsequent SCRs, DADs, and SAPs (**Fig. 6**). While RyR number does affect SCR in the cell with sparse TATS, DADs and SAPs are similarly affected in all cell types (**Fig. 6A**). Interestingly, these contrasting outcomes are similar to those reported following use of two different RyR- modulating drugs in ventricular myocytes from catecholaminergic polymorphic ventricular tachycardia (CPVT) mice. Here RyR inhibition by tetracaine decreased Ca^2+^ sparks and leak thus increasing SR Ca^2+^ load, whereas flecainide reduced Ca^2+^ spark mass and increased Ca^2+^ spark frequency with no effect on spark-mediated SR Ca^2+^ leak or content (Hilliard *et al*., 2010). Notably, changes in RyR expression may be superimposed to changes in RyR regulation, as previously discussed (Zhang et al.), which may further affect the balance between adaptive and maladaptive responses. For example, mouse knock-in RyR mutations that result in increased diastolic Ca^2+^ leak are associated with CPVT and AF, which are inhibited with the RyR stabilizer S107 that reduces diastolic SR Ca^2+^ leak in atrial myocytes and decreases burst pacing-induced AF in vivo (Shan *et al*., 2012). Additionally, AF is associated with RyR hyperphosphorylation (Vest *et al*., 2005; Neef *et al*., 2010; Voigt *et al*., 2012) and thus larger diastolic SR leak and elevated Ca^2+^ levels (Neef *et al*., 2010).

In human atrial myocytes from both normal sinus rhythm and AF patient samples, RyR density is higher at the sarcolemma than in the cell interior (Herraiz-Martínez *et al*., 2022), but the physiological significance of heterogeneous subcellular RyR distribution is not understood. In our model, increasing RyR expression at the surface vs. inner CRUs enhances NCX-RyR coupling, which on one hand reduces SCRs due to NCX-mediated Ca^2+^ extrusion, but on the other hand, increases ΔV_m_/ΔCa^2+^ gain (**Fig. 8**). Our simulations suggest that the observed increase in peripheral RyR distribution might be a protective mechanism that allows achieving the optimal balance that guarantees stable Ca^2+^ and V_m_ homeostasis. Whilst these spatial differences exist in the healthy atria, complex remodeling occurs in AF involving RyR cluster fragmentation and redistribution to inter-z-line areas, with Ca^2+^ sparks increased in fragmented CRUs (Macquaide *et al*., 2015). This subcellular remodeling of RyR distribution in AF is outside the scope of this study but can be examined in future investigations. In the companion paper (Zhang et al), we highlighted that NCX-mediated Ca^2+^ extrusion inhibits SCRs in coupled CRUs, whereas SCRs persist in uncoupled CRUs due to Ca^2+^ released having to diffuse to coupled CRUs to be extruded by NCX. In line with this, spatial differences in subcellular Ca^2+^ dynamics may be differentially affected by RyR subcellular remodeling in coupled vs. uncoupled CRUs.

### CSQ expression and distribution

CSQ is a major luminal Ca^2+^ buffer and RyR regulator (Terentyev *et al*., 2003; Györke *et al*., 2004; Knollmann *et al*., 2006; Restrepo *et al*., 2008). Reduced atrial CSQ expression promotes arrhythmia in HF (Yeh *et al*., 2008), with CSQ loss enhancing SCRs and DADs to increase AF risk (Faggioni *et al*., 2014). Our model confirms these findings and demonstrates that diminished Ca^2+^ buffering associated with lower CSQ expression has a predominant effect promoting SCRs and DADs (**Figs. 10** and **14**), with associated changes to RyR gating having the opposing (and more modest) effect of decreasing SR load to limit SCRs and DADs (**Figs. 12** and **14**). Indeed, while impaired SR Ca^2+^ buffering is a well-accepted mechanism for CPVT-associated CSQ mutations (Wleklinski *et al*., 2020), CSQ-RyR interaction may also be altered by some of these mutations (Terentyev *et al*., 2006). Notably, despite SCRs and DADs being consistently greater in the cell with sparse TATS, cells with intermediate and dense TATS are more susceptible to changes in CSQ expression (**Fig. 14**).

In human atrial myocytes, CSQ is more abundant near the cell periphery vs. the inner area (Schulson *et al*., 2011; Herraiz-Martínez *et al*., 2022). Our simulations suggest that this baseline subcellular CSQ distribution gradient might be protective against arrhythmia, as increasing surface vs. inner CRU CSQ expression promotes SCRs but inhibits DADs (**Fig. 16**). The protection from DADs is stronger in cells with sparse and intermediate TATS (**Fig. 16**), and might be relevant for chronic AF, where a cell surface to interior gradient in CSQ expression is seen and there is TATS loss (Lenaerts *et al*., 2009).

Notably, when we simulated concomitant changes in surface/inner CRU expression ratios (ratio = 2) of both RyR and CSQ to mimic experimental findings (Herraiz-Martínez *et al*., 2022), we found that the pacing threshold for SCRs increased vs varying CSQ distribution only, and varying-RyR-induced DADs are ultimately prevented by concomitant changes (i.e., CSQ + RyR) (**Fig. 19**). This indicates that appropriate relative expression of RyR and CSQ through non-uniform surface/inner CRU expression ratios, as seen in human atrial myocyte experiments (Herraiz-Martínez *et al*., 2022), provides protection against arrhythmia. Of note, adaptive changes to CSQ deficiency in CPVT include increased RyR expression, which on one hand maintains excitation-contraction coupling but also increases RyR leakiness (Song *et al*., 2007).

**Figure 19.**
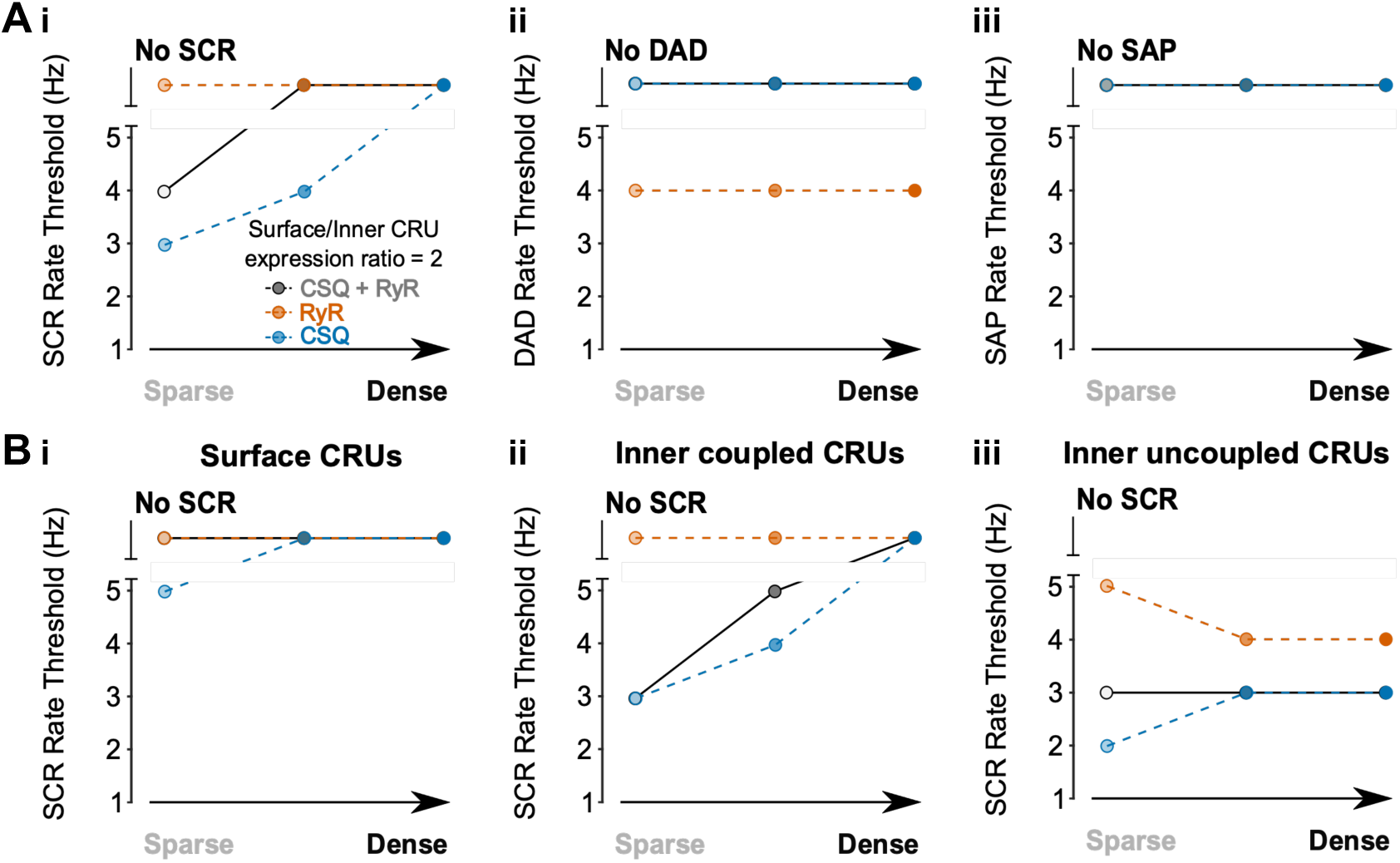
Effect of concomitantly increasing surface/inner CRU expression ratio of CSQ and RyR to 2.0. **A)** With increased CRU CSQ and RyR expression ratio, tubule loss decreases the rate threshold of SCRs (i), whereas DADs (ii) and SAPs (iii) remain absent in all conditions. The simulation results with surface/inner CRU expression ratio of RyR (orange) or CSQ (blue) at 2.0 are superimposed for comparison. **B)** Rate threshold for local SCRs from surface (i), inner coupled (ii) and inner uncoupled (iii) CRUs in sparse, intermediate, and densely tubulated cells. The impact of altered tubular density is greatest in inner coupled CRUs, with less change in SCR rate threshold associated with changes in tubule density in surface or inner uncoupled CRUs.

### LCC expression and localization

In human atrial myocytes, LCCs are evenly distributed in TATS and surface membrane (Glukhov *et al*., 2015). While LCC expression is decreased in AF (Christ T. *et al*., 2004), to our knowledge LCC subcellular distribution in AF is yet to be elucidated. In HF ventricular myocytes, altered colocalization between LCCs and RyRs may impair the ability of the LCC to trigger RyR Ca^2+^ release (Gómez *et al*., 1997). Indeed, computational atrial myocyte simulations indicate that disrupted LCC localization (i.e., removal of LCCs away from the dyadic cleft) changes subcellular Ca^2+^ dynamics to elevate SR load and enhance AP-evoked Ca^2+^ wave propagation (Shiferaw *et al*., 2020). In our preliminary simulations, we found that varying LCC expression and distribution *per se* did not change SCR and DAD properties (data not shown). Nevertheless, LCCs play a critical role in systolic Ca^2+^ signaling, as well as in setting the steady-state conditions for diastolic behavior (e.g., cellular Ca^2+^ loading and removal, which we take into account by simulating at various pacing frequencies).

## Future Directions

The limitations of this model are discussed in our companion paper (Zhang et al). Taking those into account, our novel 3D human atrial myocyte model provides a valuable platform that can be further adapted for future investigations. These could include, but are not limited to, incorporating immunofluorescence microscopy-based protein distributions and localization information of other proteins, calibrating additional experimental results of SR Ca^2+^ dynamics, and integrating extracellular ion dynamics.

### Estimation of protein localization

While spatial localization and activity of key Ca^2+^ handling proteins in the ventricle have been characterized experimentally using immunofluorescence microscopy and/or detubulation (Kawai *et al*., 1999; Despa *et al*., 2003; Gadeberg *et al*., 2017), little is currently known regarding the atria. However, various experimental data indicate heterogeneous Ca^2+^ handling proteins function, expression, and localization (Chen-Izu *et al*., 2006; Macquaide *et al*., 2015; Brandenburg *et al*., 2016; Galice *et al*., 2018; Herraiz-Martínez *et al*., 2022), and variable TATs (Trafford *et al*., 2013; Brandenburg *et al*., 2016, 2018) in atrial myocytes, which may destabilize Ca^2+^ homeostasis and the bidirectional interaction between electrical excitation and Ca^2+^ signaling. LCCs have been shown to distribute equally in tubules and crest areas of the sarcolemma (Glukhov *et al*., 2015), and rabbit atrial experiments confirm that LCCs are colocalized with RyRs as in the ventricle (Carl *et al*., 1995). Compared to the ventricle, however, atrial cells are much more diverse, with varied cell width, sarcomere spacing (Arora *et al*., 2017), and heterogeneity in the TATS between cells and across the atrium (Frisk *et al*., 2014; Gadeberg *et al*., 2016). Indeed, the subcellular distribution and expression of K^+^ ion channels, LCC, and NCX are heterogeneous in canine pulmonary veins vs. left atrium (Melnyk *et al*., 2005). While the distribution of these proteins in the healthy atria has begun to be identified, more work is needed, especially to provide insight into the remodeling that occurs in disease. Once experimental data on protein localization of human atrial myocytes becomes more readily available, this model has the potential to integrate all the spatial information to investigate how varying expression and subcellular structures may affect Ca^2+^ and voltage homeostasis.

### Effect on extracellular ion concentrations

The TATS permits rapid AP propagation but slows ion (i.e., Na^+^, Ca^2+^, and K^+^) diffusion between the extracellular space and TATS lumen (Yao *et al*., 1997; Blatter & Niggli, 1998; Shepherd & McDonough, 1998; Swift *et al*., 2006). However, myocyte contraction and relaxation cause distortion of the TATS that aids pumping the fluid to accelerate the diffusion (Savio-Galimberti *et al*., 2008; Rog-Zielinska *et al*., 2021). A computational model of Ca^2+^ diffusion within a 10-μm long tubule suggests there is a high-to-low Ca^2+^ concentration dynamic gradient between peripheral, central and deep sections, which are regulated by tubule diameters, depth, and buffering effects, extracellular ion concentration and diffusion coefficient (Pásek *et al*., 2008b). Experimental observations and mathematical models indicate the difference between tubules and surface membrane on membrane space, specific membrane capacitance and distribution of membrane ion channels, transporters and pumps in rat ventricular myocytes (Pásek *et al*., 2008a). Computational models of tubules in rat (Pásek *et al*., 2006) and guinea-pig (Pásek *et al*., 2008c) ventricular myocytes both indicate that current-clamp pacing brings transient K^+^ accumulation but Ca^2+^ depletion in the TATS lumen, whereas fast pacing weakens K^+^ accumulation but increases Ca^2+^ depletion to decrease SR load and Ca^2+^ transient amplitude. These ion concentration gradients could also be integrated into our model to investigate not only intracellular but also extracellular local Ca^2+^ dynamics.

## Conclusions

We utilized our novel 3D human atrial myocyte model to examine the arrhythmic effect of varying the expression and distribution of key Ca^2+^-handling proteins alongside changes in TATS density. We reveal a balance between the pro- and anti-arrhythmic effects of protein remodeling and that this is differentially impacted by TATS density. These findings mechanistically elucidate how remodeling underlies Ca^2+^-handling abnormalities and V_m_ instabilities that may precipitate AF.

## Competing Interests

## Abbreviations

AF: Atrial fibrillation
AP: Action potential
APD_90_: Action Potential Duration at 90% repolarization
B_CSQ_: CSQ maximum buffering capacity
CaT: Ca^2+^ transient
CPVT: Catecholaminergic polymorphic ventricular tachycardia
CRU: Ca^2+^ release unit
CSQ: Calsequestrin
DAD: Delayed afterdepolarization
I_Ca_: L-type Ca^2+^ channel current
I_K1_: Inward rectifier K^+^ current
I_K,ACh_: Acetylcholine-activated K^+^ current
I_Na_: Fast Na^+^ current
JSR: Junctional SR
LCC: L-type Ca^2+^ channel
NCX: Na^+^/Ca^2+^ exchanger
NSR: Network SR
P_O_: RyR open probability
RyR: Ryanodine receptor
SAP: Spontaneous action potential
SCR: Spontaneous Ca^2+^ release
SR: Sarcoplasmic reticulum
TATS: Transverse-axial tubular system
V_m_: Membrane voltage

